# Tractography-Pathology Correlations in Traumatic Brain Injury: A TRACK-TBI Study

**DOI:** 10.1101/2020.07.20.209668

**Authors:** Amber L. Nolan, Cathrine Petersen, Diego Iacono, Christine L. Mac Donald, Pratik Mukherjee, Andre van der Kouwe, Sonia Jain, Allison Stevens, Bram R. Diamond, Ruopeng Wang, Amy J. Markowitz, Bruce Fischl, Daniel P. Perl, Geoffrey T. Manley, C. Dirk Keene, Ramon Diaz-Arrastia, Brian L. Edlow, the TRACK-TBI Investigators

## Abstract

Diffusion tractography MRI can infer changes in network connectivity in patients with traumatic brain injury (TBI), but pathological substrates of disconnected tracts have not been well-defined due to a lack of high-resolution imaging with histopathological validation. We developed an *ex vivo* MRI protocol to analyze tract terminations at 750 μm resolution, followed by histopathologic evaluation of white matter pathology, and applied these methods to a 60-year-old man who died 26 days after TBI. Analysis of 74 cerebral hemispheric white matter regions revealed a heterogeneous distribution of tract disruptions. Associated histopathology identified variable white matter injury with patchy deposition of amyloid precursor protein and loss of neurofilament-positive axonal processes, myelin dissolution, astrogliosis, microgliosis, and perivascular hemosiderin-laden macrophages. Multiple linear regression revealed that tract disruption strongly correlated with neurofilament loss. *Ex vivo* diffusion MRI can detect tract disruptions in the human brain that reflect axonal injury.

## Introduction

Diffusion magnetic resonance imaging (dMRI) is used to study the neuroanatomic basis of altered consciousness and cognitive dysfunction in patients with traumatic brain injury (TBI).^1,2^ dMRI images can be used to map the structural connectivity of neural networks through the use of diffusion tractography, which has begun to reveal associations between brain network disconnections and their cognitive correlates.^3-8^ As the field of “connectomics” has developed, symptom mapping has transitioned from focal lesion localization to a network-based connectivity paradigm.^9,10^ For patients with TBI, this network-based approach to studying neurological deficits is particularly relevant, because disruption of neural networks by traumatic axonal injury (TAI) is one of the hallmark phenomena experienced by civilians^9^ and military personnel^11^ with head trauma.

Yet, despite the contributions that diffusion tractography and the connectomics paradigm have made to the field of TBI, fundamental questions remain about the neuropathological substrates of tract disconnections. It is well established that diffusion tractography is susceptible to false positives (i.e., anatomically implausible tracts) and false negatives (i.e., tracts that terminate prematurely where axons remain intact).^12-14^ However, observations about the limitations of diffusion tractography derive primarily from *ex vivo* MRI studies of normal, uninjured human brain specimens^15^ and non-human primate brain specimens,^16,17^ rather than analyses of human TBI brain specimens. Moreover, in the few studies that examined correlations between tract disruption and histopathology, the histological evaluation has been limited in scope, often focused only on axonal injury with reliance on amyloid precursor protein (APP) deposition to detect TAI.^18,19^ As a result, it is unknown which of the many histopathological markers of axonal injury, or associated blood brain barrier disruption and neuroinflammatory response, are associated with tract disruption in patients with TBI.

Here, we studied a human brain *ex vivo* from a patient enrolled in the longitudinal, observational <u>T</u>ransforming <u>R</u>esearch <u>a</u>nd <u>C</u>linical <u>K</u>nowledge in <u>T</u>raumatic <u>B</u>rain Injury (TRACK-TBI) study (https://tracktbi.ucsf.edu/transforming-research-and-clinical-knowledge-tbi). The patient had been admitted to the intensive care unit (ICU) and died of his injuries 26 days post-TBI. We conducted ultra-high resolution diffusion tractography and comprehensive histological analyses to determine the neuropathological correlates of tract disruption. We performed multiple linear regression to test the hypothesis that the burden of histopathological axonal injury predicts the presence of tract disruption.

## Results

### Imaging and histopathology metrics and procedures

From the brain of a 60 year-old-man who died 26 days after traumatic brain injury, we assessed pathoanatomic lesions from both *in vivo* (Supplemental Fig. 1) and *ex vivo* MRI (Supplemental Fig. 2) to facilitate comparison. We then analyzed 74 ROIs (1.5×1.5 mm^2^) throughout the cerebral hemispheric white matter, focusing on white matter pathology adjacent to the dominant left inferior frontal contusion (Fig. 1, Supplemental Fig. 3) and the grossly unremarkable white matter in the contralateral right medial and inferior frontal lobe (Supplemental Fig. 4). Within these ROIs, we quantified tract disruption using the DISCONNECT (<u>D</u>elineation of Intact and <u>S</u>evered <u>C</u>omponents <u>O</u>f <u>N</u>eural <u>NE</u>twork <u>C</u>onnec<u>T</u>ions) technique^18^ applied to the deterministic tract reconstruction from the *ex vivo* diffusion dataset (Fig. 2, Supplemental Fig. 5), and we performed quantitative histopathological analysis of axonal injury, neuroinflammation, myelin integrity, and vascular disruption (Figs. 3, 4). For tractography-pathology correlation, we assessed monotonic association between imaging and pathological metrics using Spearman’s correlation coefficient and utilized multiple linear regression to quantify how the pathologic variables in combination affected tract disruption. For a detailed description of these analytical procedures, please refer to the Methods section.

**FIG. 1.**
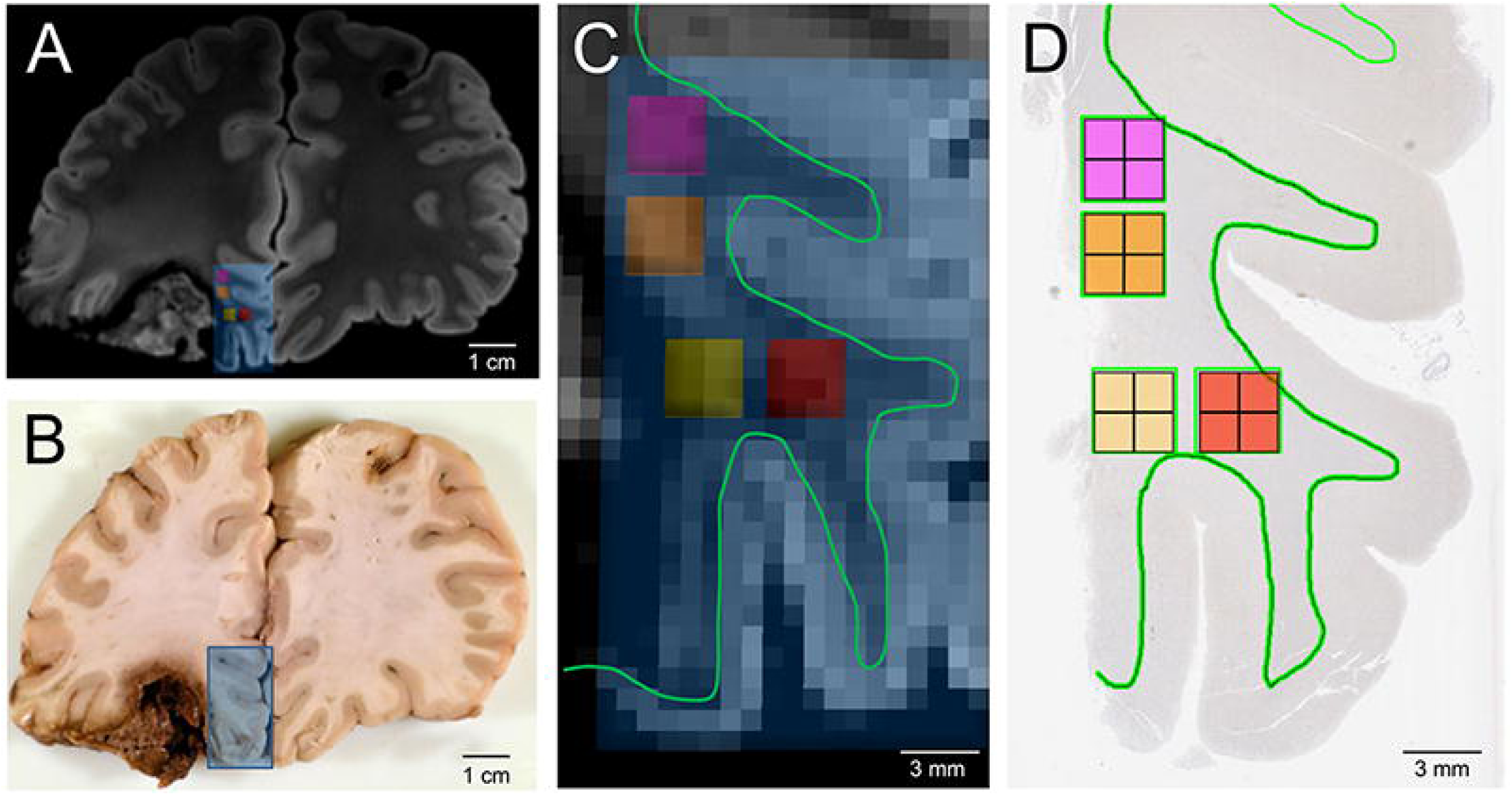
Co-registration of pericontusional regions for analysis in *ex vivo* imaging and histologic sections. A) Coronal section of MRI at the level of the left inferior frontal contusion. Blue box indicates area sampled for histology. B) Gross pathology of the same region. Blue box indicates the tissue block taken for histology. C) Magnification of area taken for histologic sampling from A. Small pink, orange, yellow and red boxes indicate regions of analysis for tractography, which were co-registered to the histology by mapping relationship to the grey white junction (green line). D) Image of tissue section with pink, orange, yellow and red boxes indicating regions of analysis for histology. A green outline of the grey-white junction is shown in (D) to match the green outline shown in (C).

**FIG. 2.**
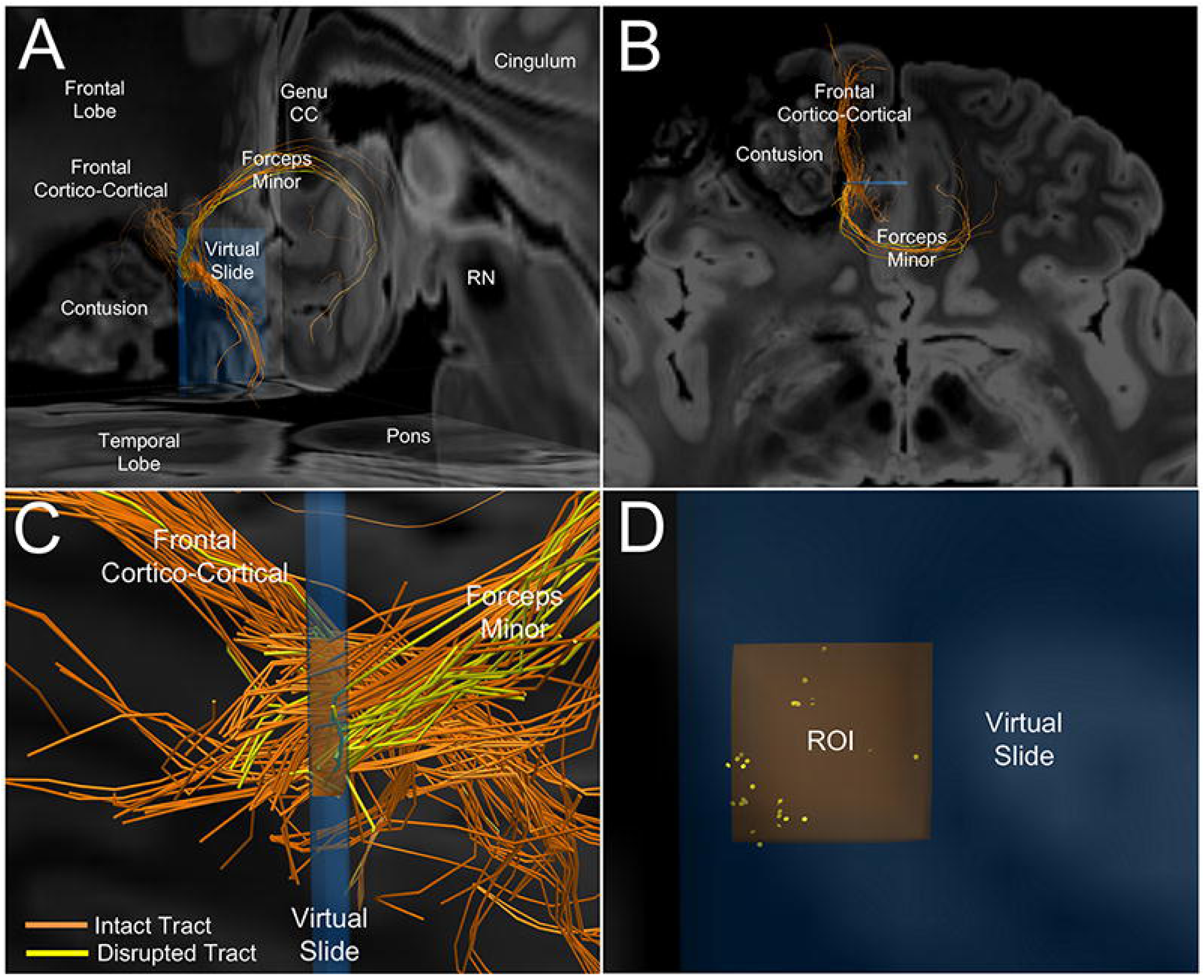
Representative image of intact and disrupted tracts passing through a pericontusional region of interest (ROI). A) Postero-lateral oblique perspective of tracts passing through the orange ROI from Figure 1. Two bundles of tracts are identified: the forceps minor crossing the genu of the corpus callosum and a frontal cortico-cortical white matter bundle. B) Higher magnification of image in A, shown from a superior perspective. Orange indicates intact tracts, whereas yellow indicates disconnected tracts that terminate in the orange ROI. C) Left lateral perspective of the orange ROI within the blue virtual slide. Intact (orange) and disconnected (yellow) tracts are seen within forceps minor and the frontal cortico-cortical bundles, with the former showing a higher percentage of disrupted tracts. D) Anterior view of disconnected tract endpoints (yellow discs) within the orange ROI reveals that the tracts tend to be disconnected in small clusters. Abbreviations: CC = corpus callosum; RN = red nucleus.

**FIG. 3.**
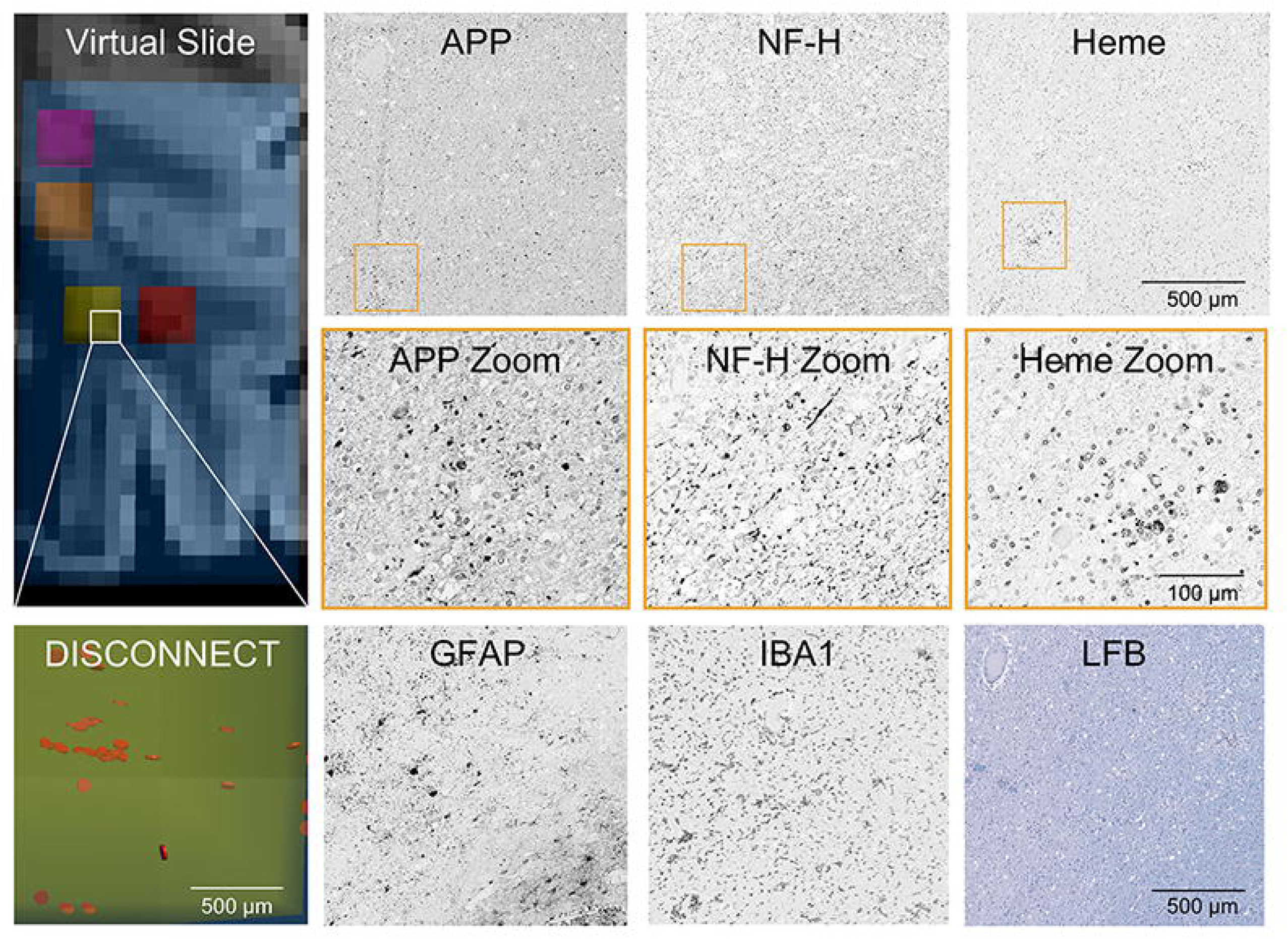
Representative histology of pericontusional region of interest with numerous disrupted tracts. The virtual slide demonstrates a pericontusional region on MRI and the associated disrupted tracts by DISCONNECT analysis; disrupted tracts are shown in orange. In the same region of interest, images are shown for immunohistochemistry of amyloid precursor protein (APP), neurofilament (NF-H), glial fibrillary acidic protein (GFAP), and ionized calcium binding adaptor molecule 1 (IBA1). Hemosiderin (Heme) deposition is identified on a counterstained slide. A Luxol fast blue (LFB) stain indicative of myelin is also displayed. Substantial deposition of APP and a lack of fine background processes with only course sparse swollen segments with NF-H is seen, consistent with the large number of disrupted tracts identified in this region.

**FIG. 4.**
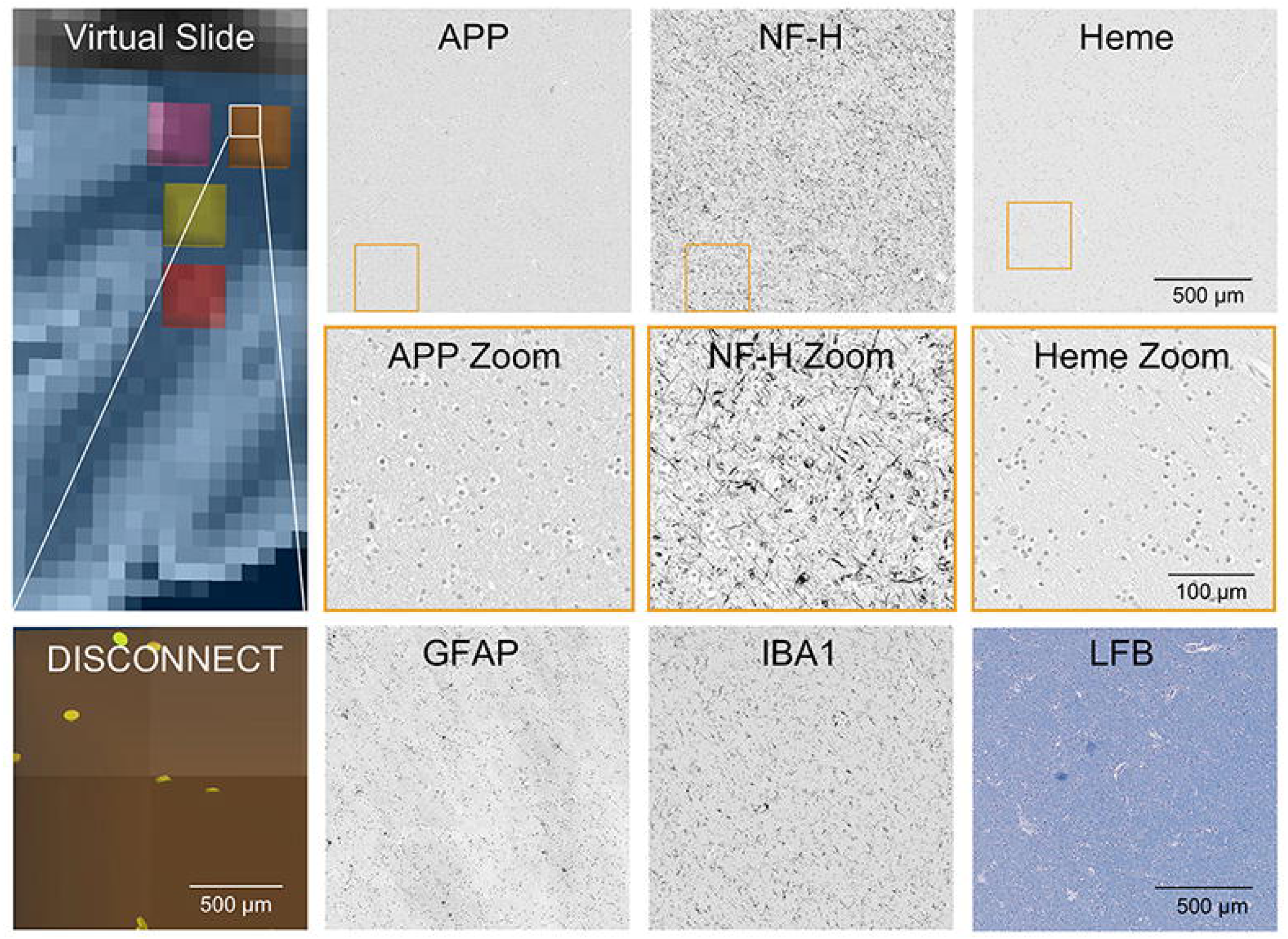
Representative histology of contralateral region of interest with few disrupted tracts. The virtual slide demonstrates a contralateral region on MRI in an area grossly intact and the associated disrupted tracts by DISCONNECT analysis; disrupted tracts are shown in yellow. In the same region of interest, images are shown for immunohistochemistry of amyloid precursor protein (APP), neurofilament (NF-H), glial fibrillary acidic protein (GFAP), and ionized calcium binding adaptor molecule 1 (IBA1). Hemosiderin (Heme) deposition is identified on a counterstained slide. A Luxol fast blue (LFB) stain indicative of myelin is also displayed. Minimal APP deposition and numerous fine background axonal processes with NF-H are seen, consistent with the small number of disrupted tracts identified in this region.

**FIG. 5.**
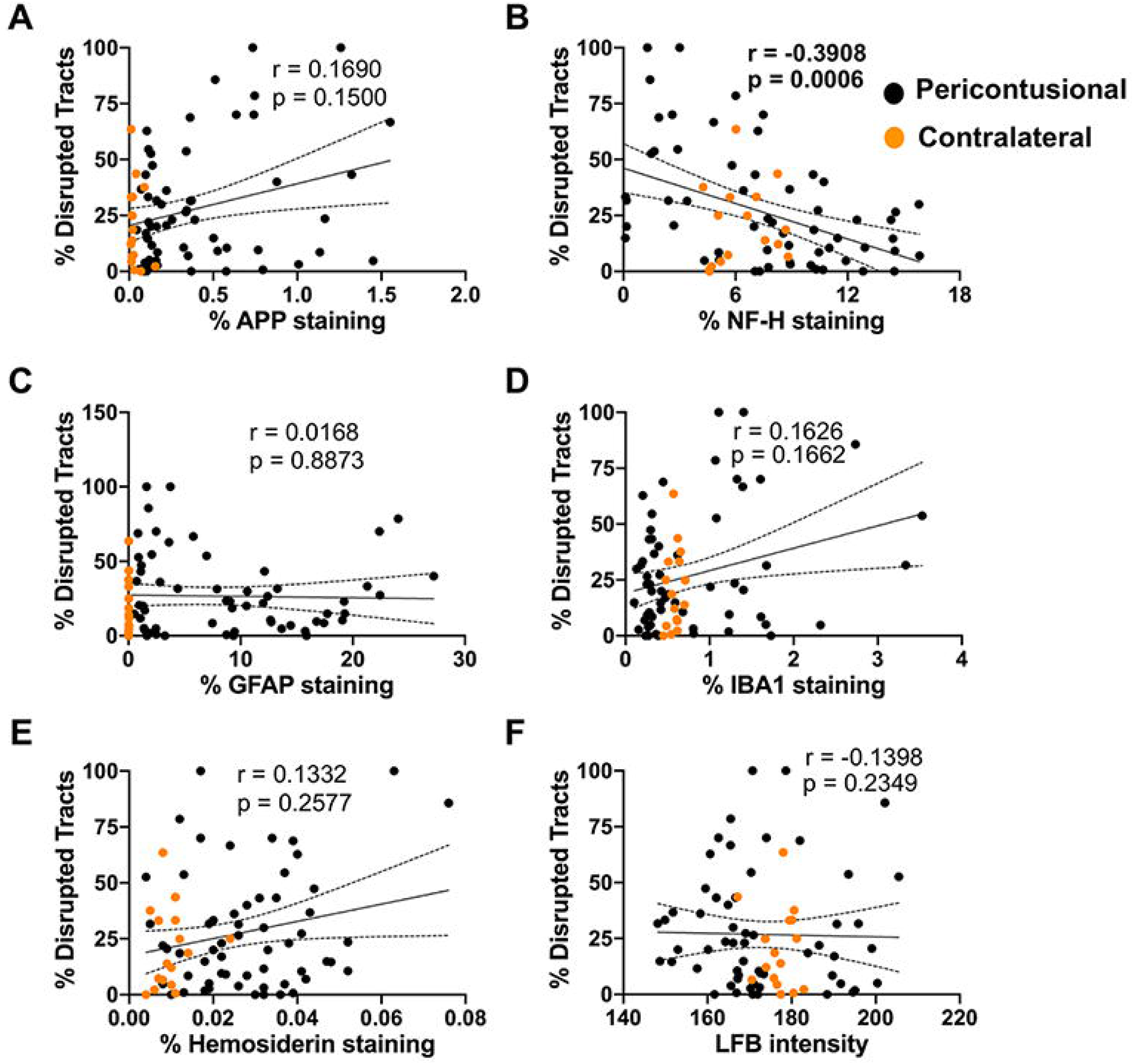
Correlations of pathology with tractography. Univariate linear regressions for each marker against % tracts disrupted are plotted: (A) amyloid precursor protein (APP), (B) neurofilament (NF-H), (C) glial fibrillary acidic protein (GFAP), (D) ionized calcium binding adaptor molecule 1(IBA1), (E) hemosiderin, and (F) Luxol fast blue (LFB). The Spearman’s correlation coefficient (r) and the associated p-value is provided for each pair. Individual data points are solid circles: black denotes pericontusional, and orange denotes contralateral regions of interest. The solid line indicates the best linear fit; dotted lines indicate the 95% confidence intervals.

### In vivo 3 Tesla MRI: Lesion Identification

Classification of the *in vivo* susceptibility weighted imaging (SWI) dataset revealed multiple pathoanatomic lesions, which were classified using the National Institutes of Health, National Institute of Neurologic Disease and Stroke’s (NINDS) Common Data Elements (CDEs) for TBI Neuroimaging (Supplemental Table 1). These data included a left frontal contusion, left temporal contusion, right frontal opercular contusion, right temporal contusion, intraventricular hemorrhage, and grade 3 diffuse axonal injury, as evidenced by traumatic microbleeds in the midbrain.

**Table 1.**
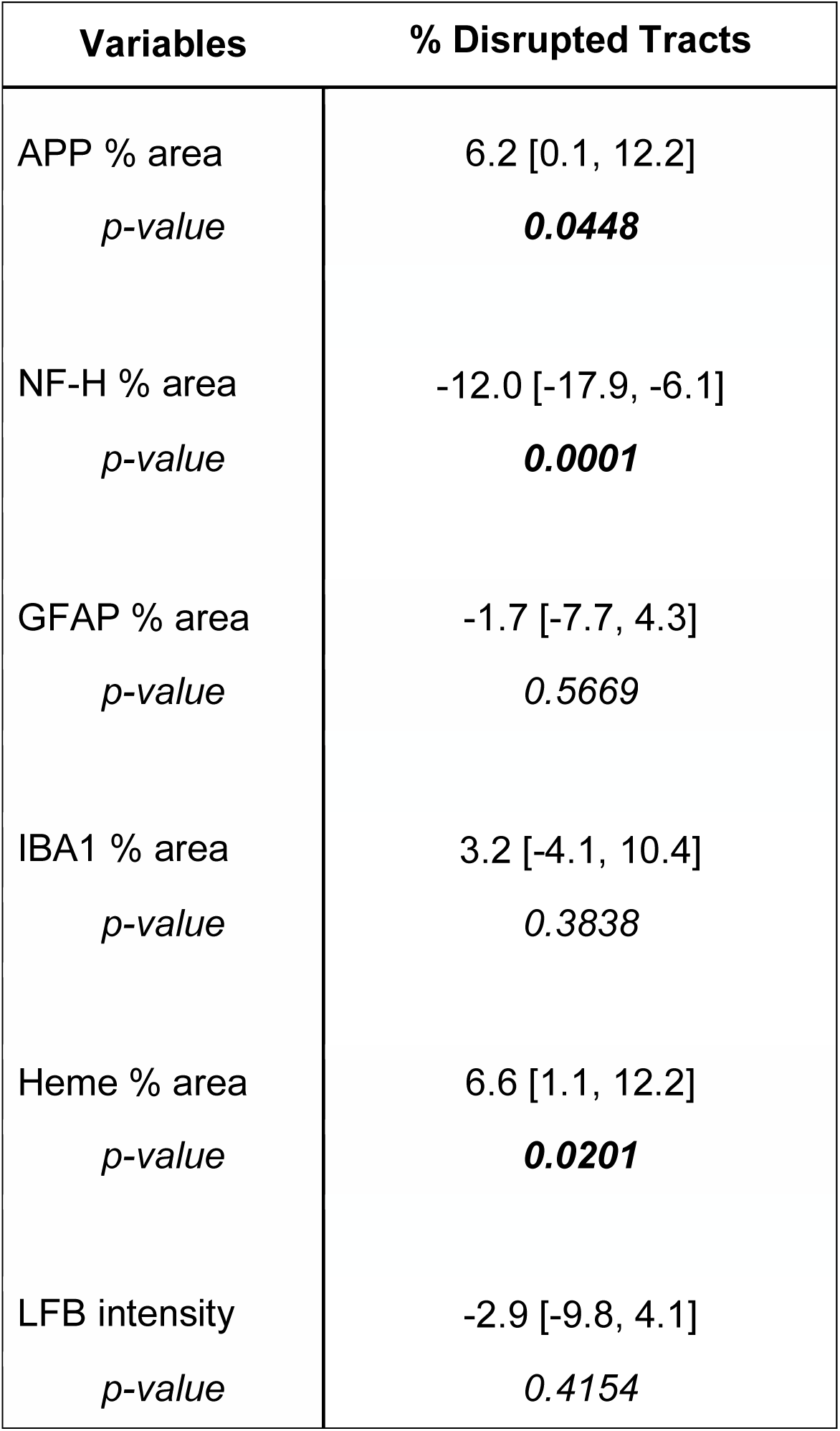
Multiple Linear Regression Analysis. Coefficient [95% Confidence Interval]; APP = amyloid precursor protein, NF-H = neurofilament, GFAP = glial fibrillary acidic protein, IBA1 = ionized calcium binding adaptor molecule 1, Heme = hemosiderin, LFB = luxol fast blue.

### Ex vivo 7 Tesla MRI: Lesion Identification

A quality assessment of the 7 Tesla MRI data revealed excellent delineation of anatomic landmarks and minimal distortions related to air bubbles (Supplemental Fig. 2). Classification of the *ex vivo* multi-echo fast low-angle shot (MEF) dataset revealed multiple lesions that were similar in signal characteristic and neuroanatomic distribution to the lesions seen on *in vivo* 3 Tesla MRI (Supplemental Table 2; Supplemental Fig. 2).

### Ex vivo 3 Tesla MRI: Tractography Disconnection Analysis

Analysis of tract termination within each of 74 cerebral hemispheric white matter ROIs using the DISCONNECT tractography technique with TrackVis software (see Methods) revealed a wide range of tract disruption. The percentage of disrupted tracts ranged from 0 to 85.7%, with a mean +/- standard deviation tract disruption of 19.9 +/- 18.0 %. These disconnections were particularly prominent in the pericontusional region, affecting multiple white matter bundles: the forceps minor (Figure 2; Supplemental Fig. 5), the cingulum bundle, the arcuate fasciculus, the medial forebrain bundle, and frontal cortico-cortical white matter bundles.

### Overview of Neuropathologic Findings

Macroscopic examination at autopsy revealed an edematous brain (formalin-fixed weight 1785 grams) with numerous contusions. The largest contusion was identified in the inferior left frontal lobe extending into the rostral left temporal lobe. The left frontal gyrus rectus remained intact, but the remainder of the left orbitofrontal and inferolateral temporal cortical surface was replaced by blood clot and granular tan-red tissue without identifiable gyral landmarks (Fig. 1, Supplemental Figs. 6, 7). A second contusion was noted in the right posterior frontal lobe localized to the inferior portion of the precentral gyrus and abutting the lateral sulcus.

Following sectioning of the brain, focal hemorrhage (∼2-3 mm) was found in the midbrain adjacent to the left red nucleus. Finally, an external ventricular drain tract was identified in the right superior frontal gyrus extending through the right centrum semiovale and right genu of the corpus callosum. Patchy accumulation of blood products was also present over the dorsal aspects of the right cerebral hemisphere, consistent with organizing subdural and subarachnoid hemorrhage. The vessels of the Circle of Willis and vertebrobasilar system were patent and intact and without significant atherosclerosis or other abnormality.

For the microscopic analysis, we focused on white matter pathology adjacent to the left inferior frontal contusion (Fig. 1, Supplemental Fig. 3), and the grossly unremarkable white matter in the contralateral right medial and inferior frontal lobe (Supplemental Fig. 4). A series of special and immunohistochemical stains were performed on serial sections to evaluate axonal damage (amyloid precursor protein [APP] accumulation and loss of phosphorylated heavy chain neurofilament [NF-H]), myelin integrity (Luxol fast blue [LFB]), reactive gliosis (glial fibrillary acidic protein [GFAP], ionized calcium binding adaptor molecule 1 [IBA1]), and hemosiderin deposition (assessed on slide with only counterstain)).

Adjacent to the left frontal contusion in a region with numerous disconnected tracts (Fig. 3), axonal pathology was prominent with frequent dot-like to globular deposition of APP. In addition, there was a loss of the usual fine fibrillar NF-H staining of axons in white matter replaced by more patchy granular deposits and occasional thickened linear fragments. An LFB stain was variable but overall pale in intensity, consistent with severe myelin loss. Frequent GFAP-positive hypertrophic astrocytes were scattered throughout the white matter and numerous reactive microglia with thickened processes and ameboid macrophages were identified with IBA1 staining. Hemosiderin-laden macrophages were often found adjacent to small capillaries.

In contrast, the grossly unremarkable white matter in the right hemisphere in a region with few disconnected tracts (Fig. 4) did not demonstrate significant APP deposition, exhibited a dense meshwork of fine NF-H-positive axonal processes, and displayed a more homogeneous and strong blue intensity with LFB staining. Scattered astrocytes with fine GFAP-positive processes were present, and IBA1 staining highlighted highly ramified microglia. Finally, significant hemosiderin deposition was not identified, overall supporting the impression of largely unremarkable white matter on histopathology and tractography. We release images of microscopic sections to the academic community at https://histopath.nmr.mgh.harvard.edu (TRACK-TBI Collection, Case ID “TRACK-NP_001”).

### Quantitative Tractography-Pathology Correlation

To examine the relationship between individual histopathology metrics and the percentage of disrupted tracts, we first assessed monotonic association using Spearman’s correlation coefficient. A negative correlation was found between NF-H staining (% area) and the percentage of disrupted tracts (Fig. 5, r = -0.39, p = 0.0006). We then built a multiple linear regression model to evaluate how the percentage of disconnected tracts was related to the pathological variables in combination. The independent variables included the percentage area of staining within the ROIs for APP, GFAP, IBA1, NF-H, and hemosiderin, as well as the intensity of LFB staining. For standardization, the independent variables were mean-centered and scaled by one standard deviation. The dependent variable was the percentage of disconnected tracts within the ROIs. In the multiple linear regression model, the percentage area of staining for NF-H, APP, and hemosiderin were the only variables to have a significant effect on the percentage of disconnected tracts (β = -12.0, p = 0.0001; β = 6.2, p = 0.0448; and β = 6.6, p = 0.0201; respectively, Table 1).

To ensure the robustness of these results, a sensitivity analysis was performed using the leave-one-out cross validation (LOOCV) method. The percentage area of NF-H staining was significantly correlated in all models with the percentage of disconnected tracts (all p < 0.001). However, a single influential ROI was identified as driving the APP correlation; when this ROI was removed from the model, a significant relationship was no longer observed between the percentage area of APP staining and the percentage of disconnected tracts (β = 3.2, p = 0.2997; Supplemental Table 3). Similar results were identified for hemosiderin staining. Two influential ROIs were noted, and when removed, a significant relationship was no longer observed between the percentage area of hemosiderin staining and the percentage of disconnected tracts (β = 4.8, p = 0.1312, Supplemental Table 4; β = 4.7, p = 0.1031, Supplemental Table 5).

## Discussion

In this correlative histo-radiologic analysis of a patient who died in the ICU from TBI, we provide persuasive evidence for a link between tract disruption and axonal injury in the human brain. Whereas prior histo-radiologic analyses of humans with TBI exposure provided qualitative evidence that tract disruption occurs in anatomic proximity to axonal injury,^18^ we implemented a quantitative approach in which ROI-based measures of tract injury were tested for correlations with gold-standard measures of white matter injury, including markers of axonal injury, demyelination, neuroinflammation, and vascular pathology. We observed that disruption of fiber tracts on *ex vivo* diffusion tractography MRI was strongly associated with loss of neurofilament staining, an expected sign of axonal loss. These findings indicate that tract disruptions reflect axonal injury in human patients with TBI.

Our findings are consistent with, and build upon, previous animal studies and limited human studies that assessed correlations between dMRI metrics and histopathological axonal injury. Similar to our findings, a model of Wallerian degeneration following dorsal root axotomy in the rat revealed reduced parallel diffusivity *in vivo* within the ipsilateral dorsal column correlating with loss of a phosphorylated heavy chain neurofilament (NF-H), over a span of 3 to 30 days after injury.^20^ In an animal model of repetitive rotational head acceleration, patterns of white matter tract damage identified by reduced fractional anisotropy were comparable to white matter injury identified by increased silver staining;^21^ likewise, silver staining reflected relative anisotropy in the corpus callosum in another mild repetitive closed head injury model.^22^ In human tissue, a correlation was identified between fractional anisotropy and axonal disruption as measured by the power coherence of myelin black gold staining at a resolution of 250 × 250 x 500 µm^3^ in cases of chronic traumatic encephalopathy.^23^

While tractography has been shown to predict cognitive and focal motor deficits in humans with severe TBI,^3,24^ few studies have directly examined correlations between tract disconnections specifically with histological axonal metrics. One study of a patient with traumatic coma evaluated spatial patterns of tract disruption with pathology, finding a similar qualitative distribution of APP immunohistochemistry and disconnected tracts near a brainstem hemorrhage.^18^ Here, we provide quantitative correlation of disconnected tracts – both nearby and remote from a focal hemorrhage – with numerous pathological indices, including measures of axonal damage and integrity, neuroinflammation, and vascular pathology.

Unexpectedly, we did not find a strong relationship between APP deposition and disconnected tracts. Only loss of neurofilament staining with an antibody to a phosphorylated heavy chain epitope (a cytoskeleton component expressed in normal white matter)^25,26^ was associated with the percentage of disrupted tracts in both monotonic correlation and multiple linear regression analyses after performing a sensitivity analysis; while the significant association of APP in the multiple linear regression model was lost after removing a single ROI. This finding might be reconciled by recent animal studies characterizing the complex temporal dynamics of axonal damage. While APP is considered the best marker for early detection of axonal injury in human tissue,^27-30^ and the degree of deposition correlates with rough groupings of TBI severity as determined by the GCS score,^31^ extensive analysis of how APP deposition changes with time, especially weeks to months after injury, has not been carefully evaluated in human tissues given inherent limitations. Animal models, however, suggest that APP alone is not sufficient to detect all types of axonal injury. For example, specific neurofilament epitopes and an N-terminal fragment of the α-spectrin protein (SNTF) accumulate in swollen axons that do not co-express APP,^32,33^ and transgenic labeling of axons in mice shows numerous axonal swellings and transections without APP accumulation after TBI.^34^

The timing and mechanism of injury also affect APP deposition. For example, at an acute time point (<24 hours after injury), Mac Donald et al. identified that anisotropy correlated with the density of APP-positive axons in a mouse model of moderate-to-severe focal contusion injury.^35^ In contrast, APP deposition has been variable or difficult to detect in tissues from less severe closed head injuries or analyzed a week to months after injury despite prominent demonstration of axonal pathology by other means.^36,37^ APP that is strongly expressed acutely after TBI in injured axons also shows decreased expression with time.^37,38^ Furthermore, APP accumulation does not always represent axonal transection, and may not indicate impending disconnection.^39^ Neurofilament loss, a direct indicator of axon loss, rather than a protein accumulation indicating changes in transport that may or may not end in permanent disconnection, may thus be a better indicator of axonal injury when correlating with tractography, especially in our patient, who died 26 days post-injury. Future studies are needed to determine how disconnected tracts correlate with different markers of axonal injury and integrity at acute to chronic time points with variable injury mechanisms.

Interestingly, indices of neuroinflammation measured with GFAP and IBA1 immunohistochemistry and myelin evaluated with LFB staining were never associated with disconnected tracts, but microvascular disruption quantified by hemosiderin deposition was a significant variable in our model when including all histological metrics and all ROIs. However, two influential, pericontusional ROIs were identified by sensitivity analysis. This observation suggests that vascular disruption can influence tractography in a specific context, such as in pericontusional locations. Vascular pathology has not been routinely included as a pathological metric in studies of dMRI radiologic-pathologic correlation. However, T2*-weighted MRI postmortem imaging has found that traumatic microbleeds defined by small foci of hypointensity represent the accumulation of perivascular iron-laden macrophages, and that this imaging metric is correlated with worse outcome in TBI.^40^ Indeed, blood brain barrier disruption and microvascular pathology has been found to be widespread even in less severe head injury and partially overlap with regions of axonal injury.^41^

Postmortem MRI has been proposed as an adjuvant to standard neuropathological assessment,^42^ as *ex vivo* MRI measures of white matter integrity may help to elucidate the neuroanatomic basis of altered consciousness and cognitive deficits observed antemortem.^18,19,43^ DISCONNECT analysis and tractography with postmortem dMRI may further highlight subtle axonal pathology not routinely examined with standard neuropathology. Use of these techniques will improve our understanding of the structural connectome of the brain even after death and fundamentally bridge how *in vivo* imaging metrics are related to complex microstructural alterations at the molecular level. Furthermore, high resolution *ex vivo* MRI methods, when performed in conjunction with gold-standard histopathological validation, may elucidate how the magnetic susceptibility effects of blood affect the reliability of tractography data in regions of hemorrhagic axonal injury.

Strengths of this study include direct quantitative correlation of numerous metrics of white matter pathology after TBI, not only axonal injury but gliosis, myelination, and vascular injury, with tract disconnections identified by ultra-high resolution tractography – an evaluation that has not been performed before in human TBI. However, several limitations should be considered when interpreting our findings. In particular, these analyses were performed in a single patient who died several weeks after injury. The pathology identified here may be multifactorial, involving pathophysiological processes such as edema and resulting local and global ischemia/hypoxia in addition to TAI. It remains to be determined whether these significant relationships are observed in TAI throughout the brain, not just pericontusional white matter, as well as other cases at variable acute, subacute and chronic time points with variable mechanisms of head injury. Furthermore, the co-registration of *ex vivo* MRI data and histopathological data was performed visually without advanced imaging techniques to correct for the known nonlinear transformations that occur with tissue processing.^44^ While the sections and ROIs used for analysis were specifically chosen to best recapitulate the relationship to the pattern of undulation of the grey-white matter junction, we cannot have achieved perfect overlap between the ROIs on imaging and pathology. Future studies are needed to optimize co-registration of *ex vivo* post-mortem MRI data with tissue sampling and analysis.

In summary, we show that axonal injury and possibly microvascular disruption may be important indices influencing diffusion tractography disconnections in TBI. Future work will address the reproducibility and nuances of these relationships in a wider patient population, providing a foundational understanding of network-based *ex vivo* imaging analyses that bridge the methodological gap between microscopy and *in vivo* MRI in human patients with TBI.

## Methods

### Case presentation

A 60-year-old man with a history of Graves’ disease and traumatic splenectomy was admitted to the hospital after a TBI. He was found down, unresponsive, next to his bicycle. An initial Glasgow Coma Scale (GCS) score of 9 (Eyes = 2, Motor = 5, Verbal = 2) was reported in the field. On admission to the emergency department, he was severely agitated and found to have a right occipital hematoma. He was sedated and intubated. Neurological examination after intubation was again notable for a GCS score of 9, with movement in response to pain in all extremities, as well as intact pupillary, corneal, cough and gag reflexes. Head CT demonstrated a right parietal skull fracture, multifocal bifrontal and temporal contusions with the largest in the left frontal lobe, a 6 mm left-sided subdural hemorrhage, diffuse convexal subarachnoid hemorrhage, and 5 mm left-to-right midline shift with transtentorial herniation. Laboratory evaluation was notable for normal complete blood cell count, electrolyte panel, and renal and coagulation parameters. The patient was immediately taken to the operating room and treated with a decompressive left hemicraniectomy and external ventricular drain placement. A repeat head CT was notable for evacuation of the subdural hemorrhage and resolution of the midline shift, as well as blossoming of the contusions, mild worsening of transtentorial herniation, and uncal herniation.

Within the first 3 days of admission to the ICU, the patient regained the ability to intermittently follow commands, and he was extubated on day 3 post-injury. However, his ICU course was complicated by a persistent fever of unknown etiology, aspiration events requiring re-intubation, focal seizure activity on electroencephalogram, a gastrointestinal bleed requiring transfusion, and progressive neurological worsening, such that he became unresponsive and developed extensor posturing in the upper extremities.

A brain 3 Tesla (3T) MRI performed on day 13 post-injury demonstrated multifocal hemorrhagic contusions in the bilateral frontal and temporal lobes, as well as subdural hemorrhage in the right frontal lobe and hemorrhagic diffuse axonal injury in the midbrain (Supplemental Fig. 1). After several weeks without neurologic improvement, the family transitioned the patient to comfort care and he died on day 26 post-injury

Prior to death, the patient was enrolled in the longitudinal, observational <u>T</u>ransforming <u>R</u>esearch <u>a</u>nd <u>C</u>linical <u>K</u>nowledge in <u>T</u>raumatic <u>B</u>rain Injury (TRACK-TBI) study (https://tracktbi.ucsf.edu/transforming-research-and-clinical-knowledge-tbi). Written informed consent for brain donation was provided by the patient’s surrogate decision-maker. Inclusion criteria for TRACK-TBI have been previously reported and include age 0-100, presentation at 1 of 18 enrolling Level 1 U.S. trauma center within 24 hours of injury with external force trauma injury to the head warranting clinical evaluation with a non-contrast head CT evaluation based on practice guidelines.^45^ Exclusion criteria include pregnancy, ongoing life-threatening disease (such as end-stage malignancy), police custody, serious psychiatric and neurologic disorders that would interfere with consent or follow-up outcome assessment, and non-English or Spanish (some sites) speakers. The study was approved by the institutional review board of each enrolling site.

### In Vivo MRI Data Acquisition and Analysis

In the *in vivo* component of this study, we focused on the SWI dataset, which provides the highest sensitivity for detection of microhemorrhages and hemorrhagic contusions.^.46^ We analyzed the SWI dataset for pathoanatomic lesions and classified each lesion using the NIH-NINDS CDEs for TBI Neuroimaging.^47^ We later applied this same lesion classification system to the *ex vivo* MRI 7 Tesla (7T) MEF MRI dataset to facilitate comparison of the *in vivo* and *ex vivo* lesion data.

### Brain specimen acquisition and fixation

The whole brain was collected with a postmortem interval of <24 hours (Supplemental Fig. 6) and suspended in 10% buffered formalin to facilitate adequate fixation, as previously described.^19^ Prior to scanning, the brain specimen was transferred from 10% buffered formalin to a Fomblin solution (perfluropolyether, Ausimont USA Inc., Thorofare, NJ) to reduce magnetic susceptibility artifacts and remove background signal.^48^ The specimen, immersed in Fomblin, was packed in a vacuum-sealed bag to minimize air bubbles that cause distortions in MRI data at the brain-air interface. Additional details regarding brain specimen packing in preparation for *ex vivo* MRI scanning have been previously described.^19^ After *ex vivo* imaging, the brain was placed back into 10% buffered formalin until brain cutting was performed.

### Ex vivo MRI acquisition

We scanned the patient’s brain using a 7 Tesla MRI scanner and 3 Tesla MRI scanner, as previously described.^19^ Briefly, the 7T Siemens Magnetom MRI scan utilized a custom-built 31-channel receive array coil^49^ and a MEF sequence^50^ at 200 µm spatial resolution with the following echo times: 5.57/ 11.77/ 17.97/ 24.17 msec. Total scan time on the 7T MRI scanner was 18 hours and 31 minutes. The 3T Siemens Tim Trio MRI scan utilized a 32-channel head coil and a 3D diffusion-weighted steady-state free-precession (DW-SSFP) sequence^51^ at 750 µm spatial resolution. The DW-SSFP sequence included 90 diffusion-weighted volumes and 12 non-diffusion-weighted volumes (*b*=0 s/mm^2^). Of note, in a DW-SSFP sequence, diffusion-weighting is not defined by a single global *b* value, because the diffusion signal cannot be readily dissociated from other imaging properties, such as the T1 relaxation time, the T2 relaxation time, TR, and the flip angle.^52^ Total diffusion scan time on the 3T MRI scanner was 30 hours and 31 minutes. All sequence parameters for the *ex vivo* 7T MEF and 3T DWSSFP sequences have been previously reported.^19^

### Ex vivo MRI processing and analysis

The 7T MEF data were processed to create tissue parameter maps,^49^ which are estimated directly from the MEF acquisitions using the DESPOT1 algorithm.^50,53^ Each parameter map provides quantification of tissue properties independent of scanner and sequence types. The parameter maps are combined to generate a synthetic FLASH scan with a flip angle of 20°, determined to have the best overall contrast and better signal-to-noise ratio than each of the contributing scans.

The 3T DWSSP data were processed using Diffusion Toolkit version 6.4.1 (http://trackvis.org/dtk) to reconstruct fiber tracts (i.e., streamlines), as previously described.^54^ Notable processing parameters in Diffusion Toolkit included: Imaging model = HARDI/Q-Ball; angle threshold = 60°; Propagation algorithm = FACT.

### MRI-guided brain cutting, tissue sampling, and immunohistopathology

Following imaging, the brainstem and cerebellum were removed, and the cerebral hemispheres were cut into ∼1 cm thick serial coronal slabs and photographed (Supplemental Fig. 7). Lesions and tissues of interest were then determined by a consensus video teleconference between investigators comparing gross morphology and anatomic landmarks on the coronal slabs with corresponding landmarks in the *ex vivo* and *in vivo* MRI. To study white matter in proximity to and remote from focal lesions, we performed tissue sampling for standard paraffin-embedded block preparation of brain parenchyma adjacent to the left inferior frontal lobe contusion, as well as in the contralateral inferior frontal lobe on the right, remote from any gross contusion or pathoanatomic lesions on imaging.

All tissue blocks were uniformly processed using an automated tissue processor (ASP 6025, Leica Biosystems, Nussloch, Germany). Standard hematoxylin and eosin (H/E) stains, as well as immunohistochemical and special stains, were performed on serial 5 µm thick sections. Special stains included LFB. Immunohistochemical stains included antibodies against APP, GFAP, IBA1, and neurofilament, specifically to NF-H, which shows expression in uninjured axons.^25^ All immunostains were performed using a Leica Bond III automated immunostainer with a diaminobenzidine chromogen detection system (DS9800, Leica Biosystems, Buffalo Grove, IL, USA).

Specifications for each antibody are as follows: anti-amyloid precursor protein (APP; mouse anti-human monoclonal antibody clone 22c11, dilution 1:10, epitope retrieval time 10 min, MAB348; EMD Millipore, Burlington, MA, USA); anti-glial fibrillary acidic protein (GFAP; mouse anti-human monoclonal antibody GA5, with bond heat-induced epitope retrieval, epitope retrieval time 10 min, PA0026; Leica Biosystems, Wetzlar, Germany); anti-ionized calcium-binding adapter molecule 1 (IBA1; rabbit polyclonal, dilution 1:100, epitope retrieval time 10 min, Wako 016-20001; FUJIFILM Wako Pure Chemical Corporation, Osaka, Japan); anti-neurofilament to a phosphorylated heavy chain neurofilament (NF-H; mouse anti-human monoclonal antibody SMI-34, dilution 1:100, epitope retrieval time 10 min, Biolegend 835503; Biolegend, San Diego, CA). Hemosiderin deposition was evaluated on a section with only counterstain. All stained slides were scanned at 20x magnification into digital images using the Aperio scanner system (Aperio AT2 - High Volume, Digital whole slide scanning scanner, Leica Biosystems Inc., Richmond, IL, USA) for analysis.

### Region of Interest Selection for Histo-Radiologic Correlation Analysis

To optimize the spatial alignment of the histopathological and radiological data, we rotated the coronal plane of the diffusion dataset using FreeView software (https://surfer.nmr.mgh.harvard.edu/fswiki/FreeviewGuide) (Fig. 1, Supplemental Figs. 3, 4). Visual delineation of corresponding anatomic landmarks was performed by a neuropathologist (A.N.), critical care neurologist (B.L.E.) and research technician (B.R.D.). Once an optimal rotation was agreed upon, this rotation of the diffusion dataset was applied to the tractography data in TrackVis, allowing all tractography analyses to be performed in a spatial coordinate system that closely matched that of the coronal gross pathology slabs.

Next, we created “virtual slides” in TrackVis with the same dimensions as the sampled tissue sections (Fig. 1, Supplemental Figs. 3, 4). Within each pair of virtual slides and histopathology slides, we created ROIs for correlative tractography-histopathology analysis. These square-shaped ROIs were matched through visual inspection, based upon the curvature of the grey-white junction. We placed the ROIs within the white matter, consistent with this study’s aim of detecting the histopathological signatures of tract disruptions. Although directional water diffusion may be present within the grey matter of the cerebral cortex,^55^ tracts typically terminate when they reach normal grey matter, and thus grey matter was not probed in this study.

We then considered the optimal ROI size for testing correlations between tract disruptions and histopathological markers of axonal white matter injury. We used a data-driven approach in which we analyzed clusters of disrupted tracts in white matter pathways that were not included in the virtual slides: the genu, body, and splenium of the corpus callosum (Supplemental Figs. 8, 9). Visual analysis of disrupted tracts within these white matter bundles demonstrated that tract disruptions were typically localized to square-shaped clusters of four voxels in a 2×2 grid (1.5 mm x 1.5 mm) or smaller (Supplemental Figs. 8, 9). We therefore created 4-voxel, square-shaped ROIs for testing tractography-pathology correlations in the virtual slides. Selection of larger ROIs would have lowered our statistical power to detect an association between tract disruption and histopathological signs of axonal injury, because intact tracts would have been interspersed with disconnected tracts within each ROI. Of note, groups of 4 ROIs were spatially arranged in a square, which allowed us to optimize visual co-registration between the histopathology and tractography data and minimize the number of anatomic landmarks required to identify each ROI (Fig. 1, Supplemental Figs. 3, 4). We first assessed 64 ROIs, of which most were placed >2 mm from the left inferior frontal contusion. However, we observed higher levels of tract disruption and prominent pathology in a few ROIs directly adjacent to the contusion. Given that this pathology is of biological significance, we believed that it should be well represented in our model, and thus we added 12 ROIs directly adjacent to the contusion (brown, cream and light pink boxes in Supplemental Fig. 3). ROIs were excluded if no tracts were detected. 76 ROIs were evaluated in total, and 2 were excluded.

### Tractography Analysis

Once ROI size and placement was determined within the virtual slides, we processed the *ex vivo* diffusion dataset for deterministic tract construction using Diffusion Toolkit, and we performed a tractography analysis: DISCONNECT using the TrackVis software (www.trackvis.org).^18^ The DISCONNECT technique allows for a virtual dissection of disrupted fiber tracts from intact fiber tracts, with visualization and quantification of tract “end-points” that terminate within an ROI. Given that tracts are not expected to terminate within healthy white matter, the presence of a tract endpoint is interpreted as inferential evidence of axonal injury, leading to a loss of directional water diffusion along the injured axonal bundle. For this study, TrackVis creator Ruopeng Wang, MS, created new functionality in TrackVis that allows focused visualization of tract endpoints within an ROI (i.e., without concurrently visualizing the other end of the tract). We release this new DISCONNECT functionality in an updated version of the TrackVis software program as version 0.6.3 (www.trackvis.org).

### Histopathology Analysis

To quantify the white matter pathology and allow correlation with tractography, several metrics were extracted from scanned images of the immunohistochemical and special stains. The staining was quantified for APP, NF-H, GFAP, IBA1 and hemosiderin as follows: Aperio ImageScope^©^ (Aperio ImageScope, version 2016, Leica Biosystems Inc., Richmond, IL, USA) was used to extract a tiff image containing 4 ROIs (each ROI 1.5×1.5 mm^2^). All images were then processed and quantified using FIJI/ImageJ software.^56^ Each image was divided into 4 (for final ROIs), converted into greyscale and then a binary image, using the same predetermined threshold for every image. Expression or deposition was quantified as the percentage of area staining. The LFB-stained slide was quantified as the average intensity of staining on an RGB image rather than the percent area after thresholding. In some ROIs, directly adjacent to the contusion edge, a small area without tissue was included within the 1.5×1.5 mm area. These “blank” areas were manually cropped from the image prior to analysis, and percentage of area or intensity of staining was only calculated within the tissue present.

### Statistical Analysis

Statistical analyses were performed using R Statistical Software (version 3.6.1; R Foundation for Statistical Computing, Vienna, Austria). Univariate monotonic correlation between variables was analyzed using Spearman’s correlation coefficient. Multiple linear regression evaluated the relationship between the percentage of disconnected tracts and the standardized quantitative pathological data, including the percentage of staining for APP, GFAP, IBA1, NF-H, and hemosiderin and the intensity of LFB. The predictor variables were mean-centered and scaled by one standard deviation. These quantitative pathological variables were selected as the best indicators of the relative presence of each marker within each ROI. The percentage of disconnected tracts rather than the number of disconnected tracts was chosen as the outcome variable given the variability in tract number detected within the ROIs. Multicollinearity among the predictors was investigated by using Spearman’s correlation and the variance inflation factor. Sensitivity analysis was performed on the multiple linear regression model using the leave-one-out cross validation (LOOCV) method whereby one observation was systematically removed from the model to evaluate the effect on the significance of the contributing variables. P-values < 0.05 were considered statistically significant.

## Supporting information

FinalDataSet

## Data Availability

The dataset used for final analysis is provided as a supplemental excel file. Histopathological images are available at https://histopath.nmr.mgh.harvard.edu (TRACK-TBI Collection, Case ID “TRACK-NP_001”). The DISCONNECT functionality for tract termination visualization is available in an updated version of the TrackVis software program as version 0.6.3 (www.trackvis.org). Additional data requests can be made to Brian L. Edlow, M.D.

## Acknowledgments

We thank the patient and family for their participation in the TRACK-TBI study and for their generous donation of the post-mortem brain. The study was funded by the NIH National Institute of Neurological Disorders and Stroke (U01NS086090, K08NS114170, R21NS109627, RF1NS115268, K23NS094538, R01NS0525851, R21NS072652, R01NS070963, R01NS083534, U01NS086625, U24NS10059103, R01NS105820), NIH Director’s Office (DP2HD101400), NIH National Institute for Biomedical Imaging and Bioengineering (P41EB015896, 1R01EB023281, R01EB006758, R21EB018907, R01EB019956), NIH National Institute on Aging (R56AG064027, R01AG064027, R01AG008122, R01AG016495), NIH National Institute of Diabetes and Digestive and Kidney Diseases (R21DK108277), James S. McDonnell Foundation, Tiny Blue Dot Foundation, and the Nancy and Buster Alvord endowment (CDK). This research also utilized resources provided by NIH shared instrumentation grants S10RR023401, S10RR019307, and S10RR023043. Additional support was provided by the BRAIN Initiative Cell Census Network (U01MH117023) and the NIH Blueprint for Neuroscience Research (5U01-MH093765), part of the multi-institutional Human Connectome Project, and One Mind.

## Author Contributions

A.L.N and B.L.E conceptualized the study. A.L.N analyzed the histopathology data. B.L.E analyzed the tractography data. C.P. performed the statistical analysis and modeling. S.J. provided critical input for the statistical analysis. A.L.N and B.L.E wrote the manuscript. D.I. and D.P.P assisted with the neuropathology characterization, tissue sampling, immunohistochemical staining and slide scanning. C.L.M and P.M assisted with radiological-pathological correlation for tissue sampling and provided methodological input. A.v.d.K., A.S., B.R.D, R.W. and B.F. assisted with ex vivo imaging acquisition, processing, tractography analysis, and methodological input. A.J.M edited the manuscript. G.T.M., C.D.K., and R.D-A. provided methodological input and interpretation of results. All authors revised the manuscript and provided critical feedback.

