## Supplemental Material for "Tractography-Pathology Correlations in Traumatic Brain Injury: A TRACK-TBI Study"

| CDE | Present? | Laterality | Anatomic Site | Comments |
|---|---|---|---|---|
| Skull fracture | Y | Right | Parietal | Also with skull defect due to left hemicraniectomy |
| Epidural hemorrhage | N | N/A | N/A | N/A |
| Subdural hemorrhage | Y | Bilateral | Fronto-temporo-parietal | N/A |
| Subarachnoid hemorrhage | Y | Right | Sylvian fissure and frontoparietal sulci | Sylvian fissure subarachnoid hemorrhage is adjacent to small right fronto-temporal contusions |
| Contusion | Y | Bilateral | Bifrontal, bitemporal | Left frontal and left temporal contusions larger than right frontal and right temporal contusions |
| Traumatic Axonal Injury | Y | bilateral | Hemispheric white matter, as well as genu of the corpus callosum, bilateral midbrain tegmentum, and ventral pons | N/A |
| Edema | Y | Left | Pericontusional edema within the left frontal and left temporal contusions | N/A |
| Ischemic or hypoxic-ischemic injury | N | N/A | N/A | N/A |
| Brain atrophy or encephalomalacia | N | N/A | N/A | N/A |

**Supplemental Table 1: Traumatic Lesions Visualized on *In Vivo* MRI.**
All pathoanatomic lesions are characterized using the NIH Common Data Element (CDE) Guidelines for Traumatic Brain Injury Neuroimaging. N/A = not applicable.

| CDE | Present? | Laterality | Anatomic Site | Comments |
|---|---|---|---|---|
| Skull fracture | Unable to Assess | Unable to Assess | Unable to Assess | Unable to Assess |
| Epidural hemorrhage | Unable to Assess | Unable to Assess | Unable to Assess | Unable to Assess |
| Subdural hemorrhage | Unable to Assess | Unable to Assess | Unable to Assess | Unable to Assess |
| Subarachnoid hemorrhage | Y | Right | Sylvian fissure | Adjacent to small right fronto-temporal contusions |
| Contusion | Y | Bilateral | Bifrontal, bitemporal | Left frontal and left temporal contusions larger than right frontal and right temporal contusions |
| Traumatic Axonal Injury | Y | bilateral | Hemispheric white matter, as well as genu of the corpus callosum, bilateral midbrain tegmentum, and ventral pons | N/A |
| Edema | Y | Left | Pericontusional edema within the left frontal and left temporal contusions | N/A |
| Ischemic or hypoxic-ischemic injury | N | N/A | N/A | N/A |
| Brain atrophy or encephalomalacia | N | N/A | N/A | N/A |

**Supplemental Table 2: Traumatic Lesions Visualized on *Ex Vivo* MRI.**
All pathoanatomic lesions are characterized using the NIH Common Data Element (CDE) Guidelines for Traumatic Brain Injury Neuroimaging. N/A = not applicable.

| Variables | % Disrupted Tracts |
|---|---|
| APP % area | 3.2 (-2.9, 9.2) |
| *p-value* | *0.2997* |
| NF-H % area | -11.6 (-17.3, -5.9) |
| *p-value* | ***0.0001*** |
| GFAP % area | -0.2 (-6.1, 5.8) |
| *p-value* | *0.9548* |
| IBA1 % area | 2.9 (-4.1, 9.8) |
| *p-value* | *0.4172* |
| Heme % area | 7.9 (2.4, 13.4) |
| *p-value* | ***0.0052*** |
| LFB intensity | -1.9 (-8.7, 4.9) |
| *p-value* | *0.5854* |

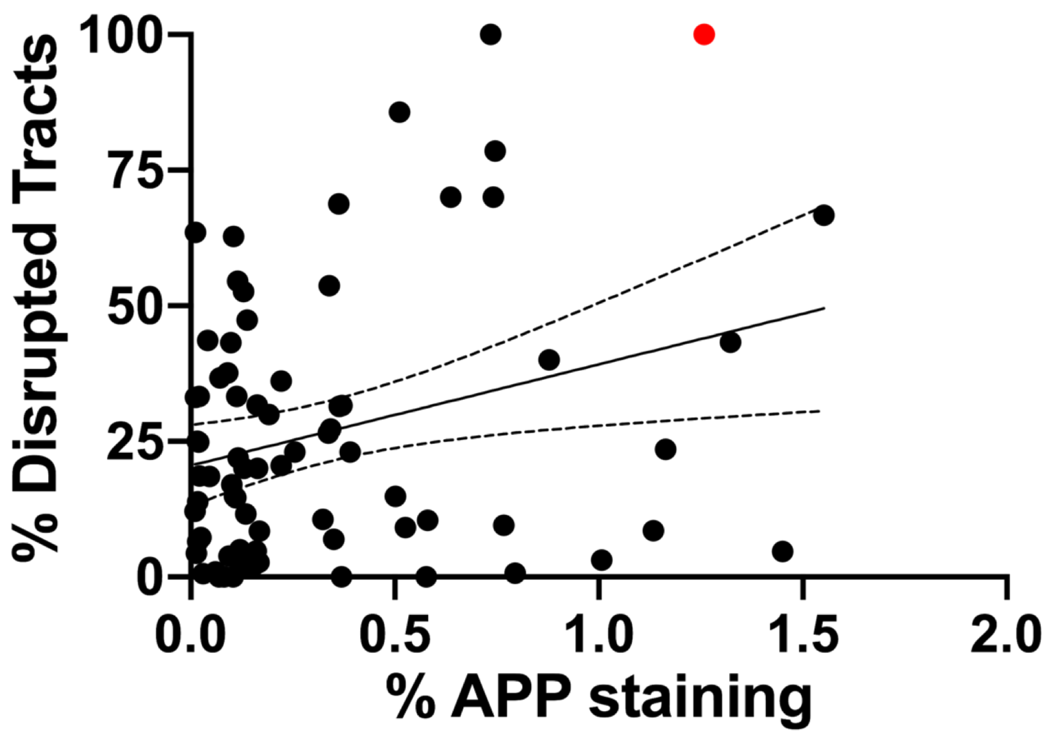


**Supplemental Table 3. Multiple Linear Regression Analysis without single ROI driving APP correlation indicated in red.** Coefficient (Confidence Interval); APP = amyloid precursor protein, NF-H = neurofilament, GFAP = glial fibrillary acidic protein, IBA1 = ionized calcium binding adaptor molecule 1, Heme = hemosiderin, LFB = Luxol fast blue.

| Variables | % Disrupted Tracts |
|---|---|
| APP % area | 6.6 (0.4, 12.7) |
| *p-value* | ***0.0366*** |
| NF-H % area | -11.5 (-17.4, -5.5) |
| *p-value* | ***0.0003*** |
| GFAP % area | -1.8 (-7.9, 4.2) |
| *p-value* | *0.5488* |
| IBA1 % area | 2.6 (-4.4, 9.6) |
| *p-value* | *0.4587* |
| Heme % area | 4.8 (-1.5, 11.0) |
| *p-value* | *0.1312* |
| LFB intensity | -3.4 (-10.5, 3.6) |
| *p-value* | *0.3307* |

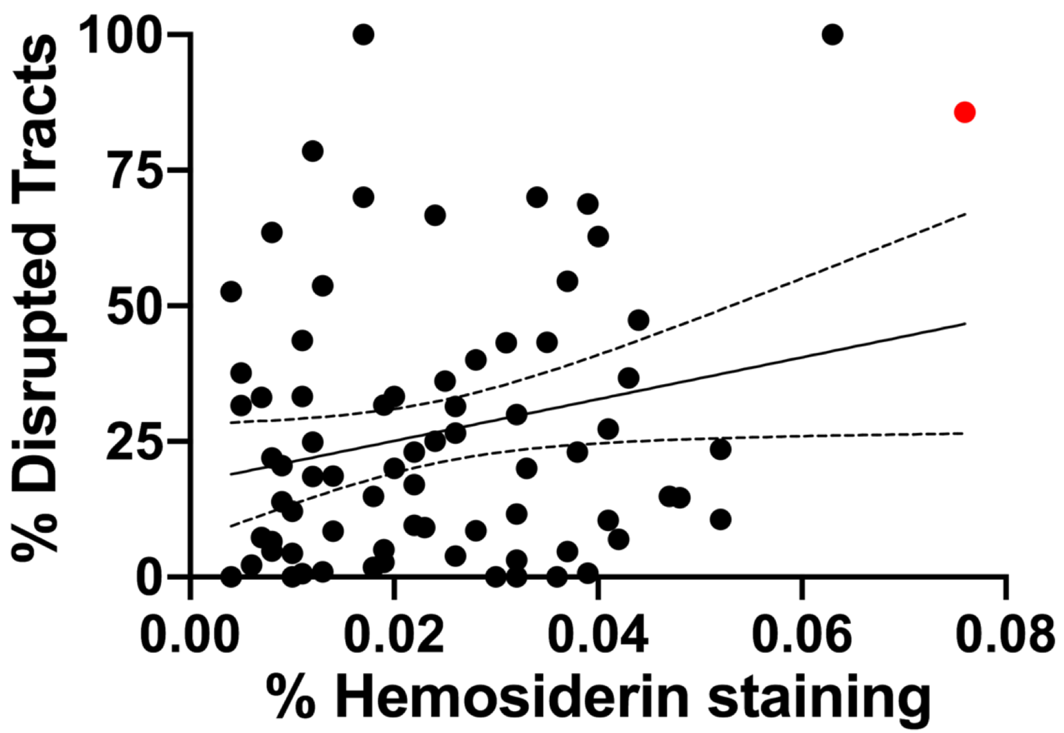


**Supplemental Table 4. Multiple Linear Regression Analysis without one or two ROIs driving Heme correlation indicated in red.** Coefficient (Confidence Interval); APP = amyloid precursor protein, NF-H = neurofilament, GFAP = glial fibrillary acidic protein, IBA1 = ionized calcium binding adaptor molecule 1, Heme = hemosiderin, LFB = Luxol fast blue.

| Variables | % Disrupted Tracts |
|---|---|
| APP % area | 5.7 (-0.3, 11.7) |
| *p-value* | *0.0608* |
| NF-H % area | -10.6 (-16.6, -4.6) |
| *p-value* | ***0.0007*** |
| GFAP % area | -1.4 (-7.3, 4.6) |
| *p-value* | *0.6516* |
| IBA1 % area | 4.0 (-3.3, 11.2) |
| *p-value* | *0.2795* |
| Heme % area | 4.7 (-1.0, 10.3) |
| *p-value* | *0.1031* |
| LFB intensity | -3.7 (-10.7, 3.2) |
| *p-value* | *0.2893* |

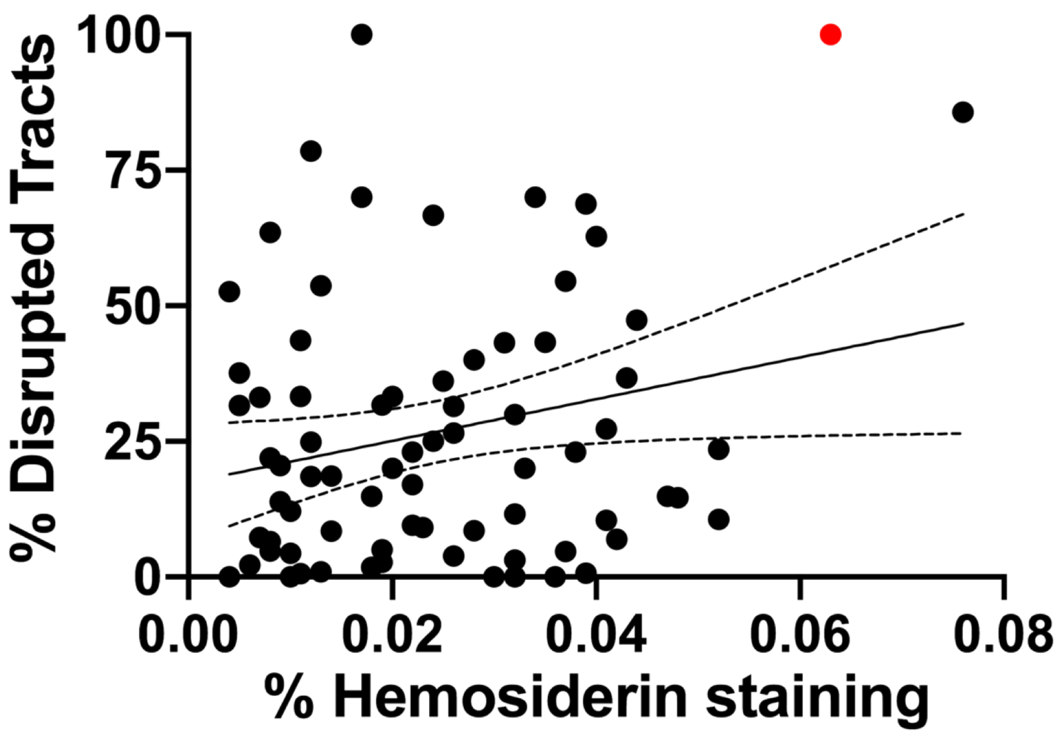


**Supplemental Table 5. Multiple Linear Regression Analysis without one or two ROIs driving Heme correlation indicated in red.** Coefficient (Confidence Interval); APP = amyloid precursor protein, NF-H = neurofilament, GFAP = glial fibrillary acidic protein, IBA1 = ionized calcium binding adaptor molecule 1, Heme = hemosiderin, LFB =Luxol fast blue.

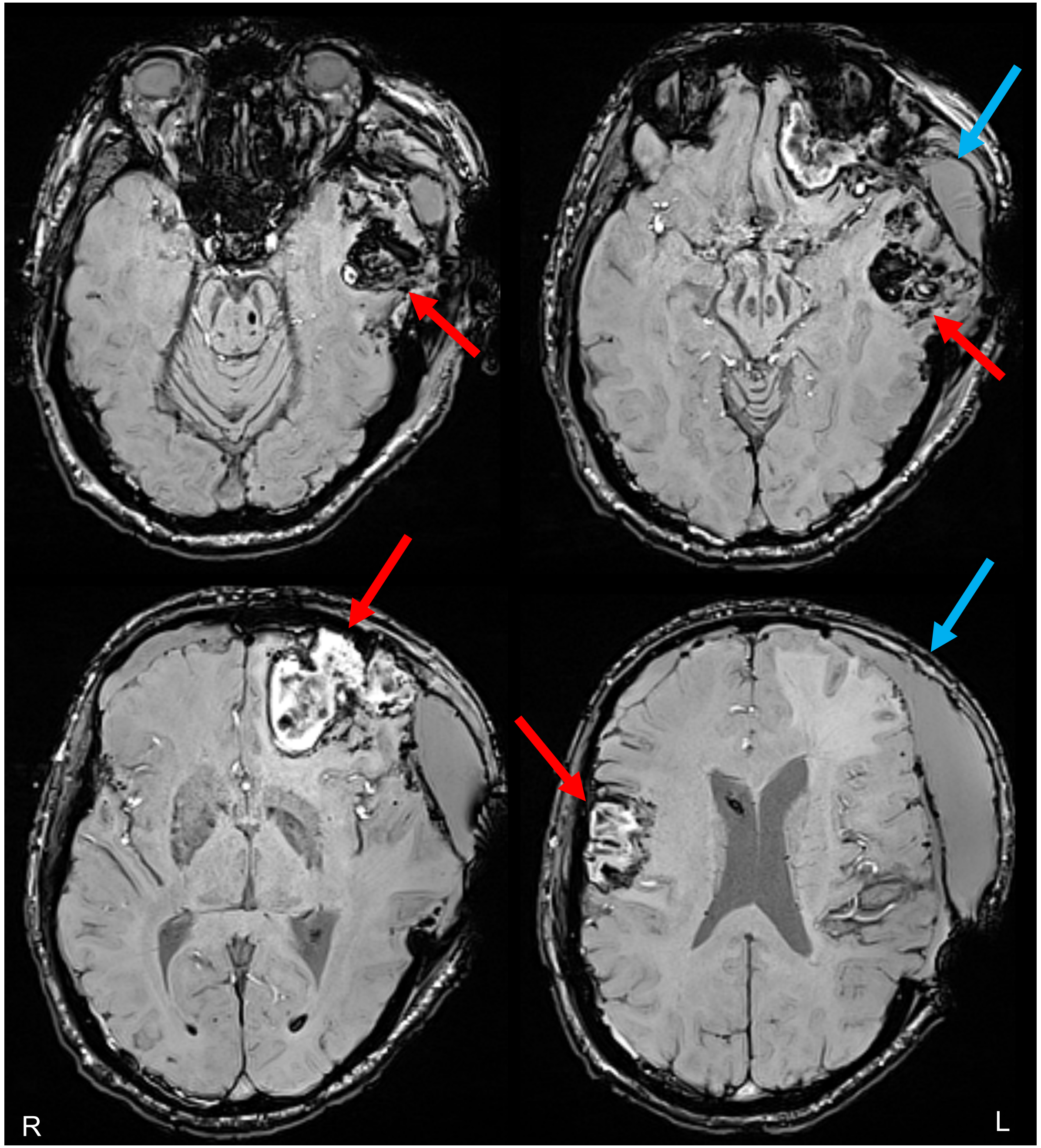


**Supplemental Figure 1**. **Susceptibility-weighted *in vivo* MRI performed on day 13 post-injury.** Axial images are shown at the level of the caudal midbrain (top left), rostral midbrain (top right), mid-thalamus (bottom left), and septum pellucidum (bottom right). Multifocal contusions are indicated by red arrows in the left frontal lobe, left temporal lobe, and right frontal lobe. An extra-axial fluid collection is also seen in the left frontotemporal region (blue arrows), at the site of the recent left hemicraniectomy. In the top right image, the medial temporal lobes are seen compressing the dorsal aspect of the rostral midbrain bilaterally. R = right, L = left.

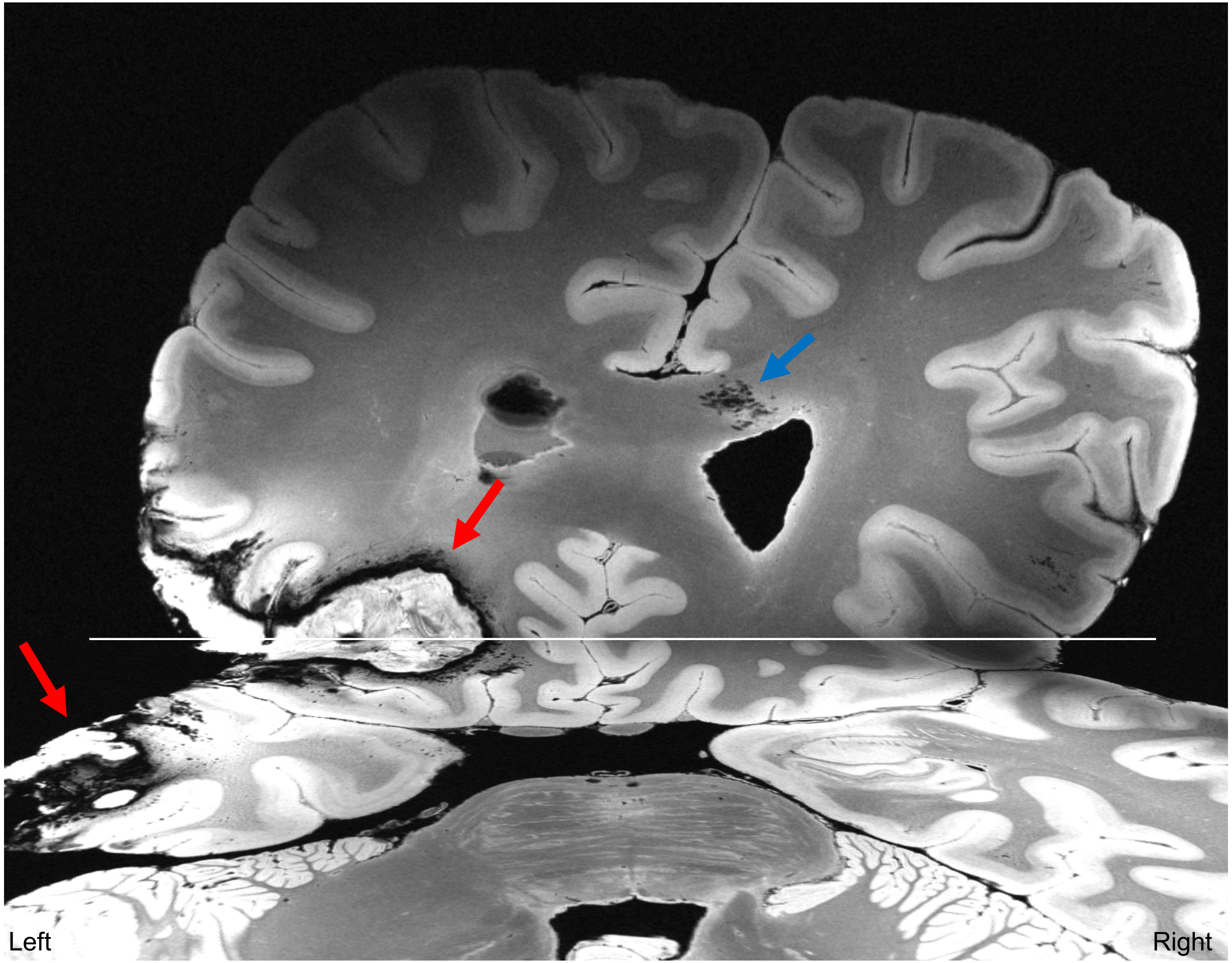


**Supplemental Figure 2. Quality Assessment of *ex vivo* 7 Tesla Multi-echo Flash (MEF) Dataset.** A posterior view of an axial slice at the level of the mid-pons (foreground) is shown, along with a coronal section in the plane of the left frontal contusion. The horizontal white line indicates the intersection of the axial and coronal planes. Both images are from a MEF flash parameter map (flash20) at 200 μm spatial resolution. The MEF images provide clear anatomic delineation of cortical and subcortical structures, as well as identification of the left frontal and temporal contusions (red arrows). Punctate foci of hemorrhage, likely representing hemorrhagic traumatic axonal injury, are also seen in the genu of the corpus callosum (blue arrow). Of note, the images shown here are from a combined synthetic FLASH scan with a flip angle of 20˚.

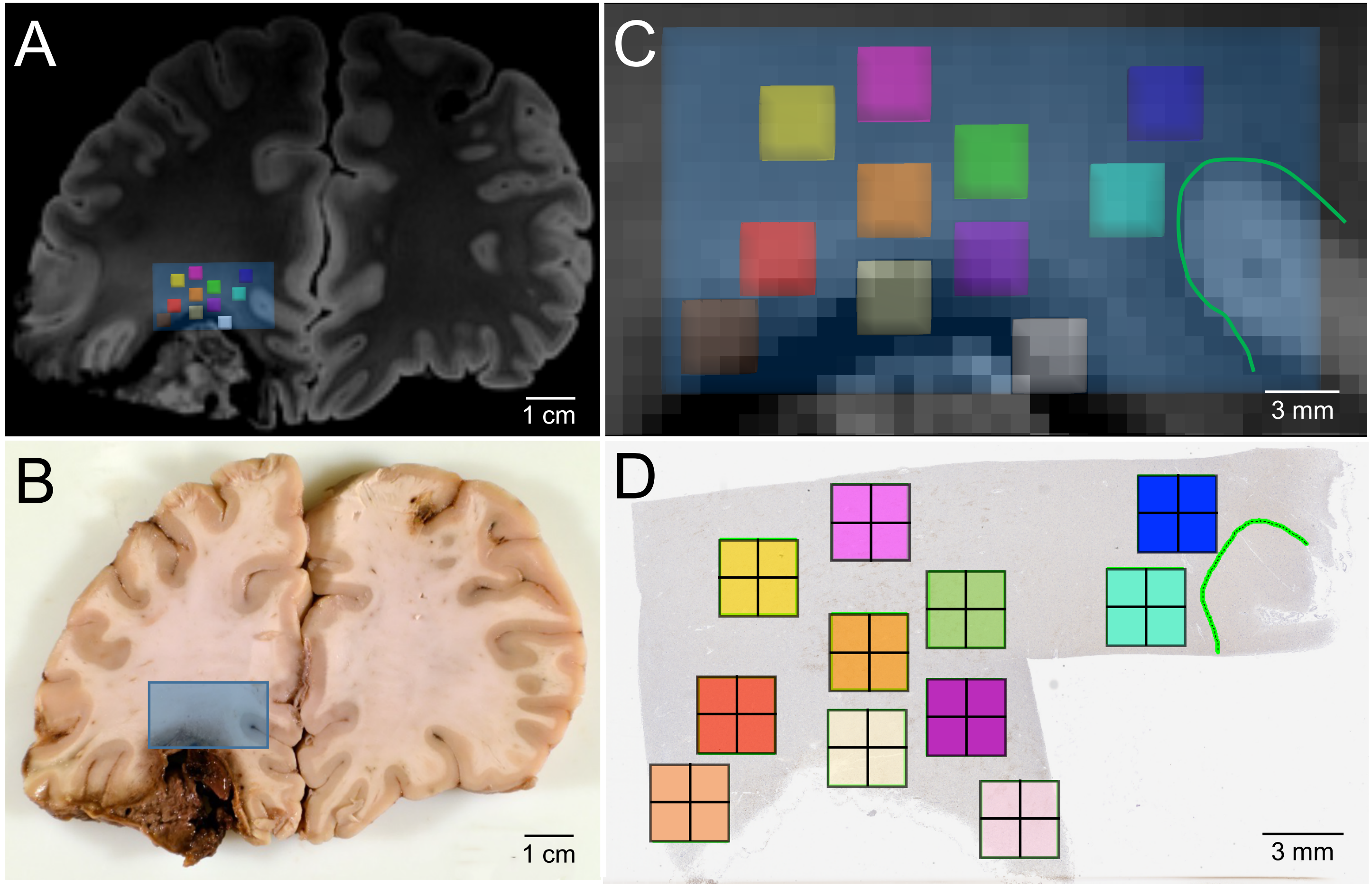


**Supplemental Figure 3. Coregistration of pericontusional regions for analysis in *ex vivo* imaging and histologic sections.**
A) Coronal section of MRI at the level of the left inferior frontal contusion. Blue box indicates area sampled for histology. B) Gross pathology of the same region. Blue box indicates the tissue block taken for histology. C) Magnification of area taken for histologic sampling from A. Small yellow, red, pink, orange, green, purple, turquoise, dark blue, brown, cream and light pink boxes indicate regions of analysis for tractography, which were co-registered to the histology by mapping their anatomic relationship to the grey-white junction. The green outline in (C) indicates the grey-white junction at the depth of the cortical sulcus. D) Image of tissue section with matching color-coded boxes, indicating regions of analysis for histology. A green outline of the grey-white junction is shown in (D) to match the green outline shown in (C).

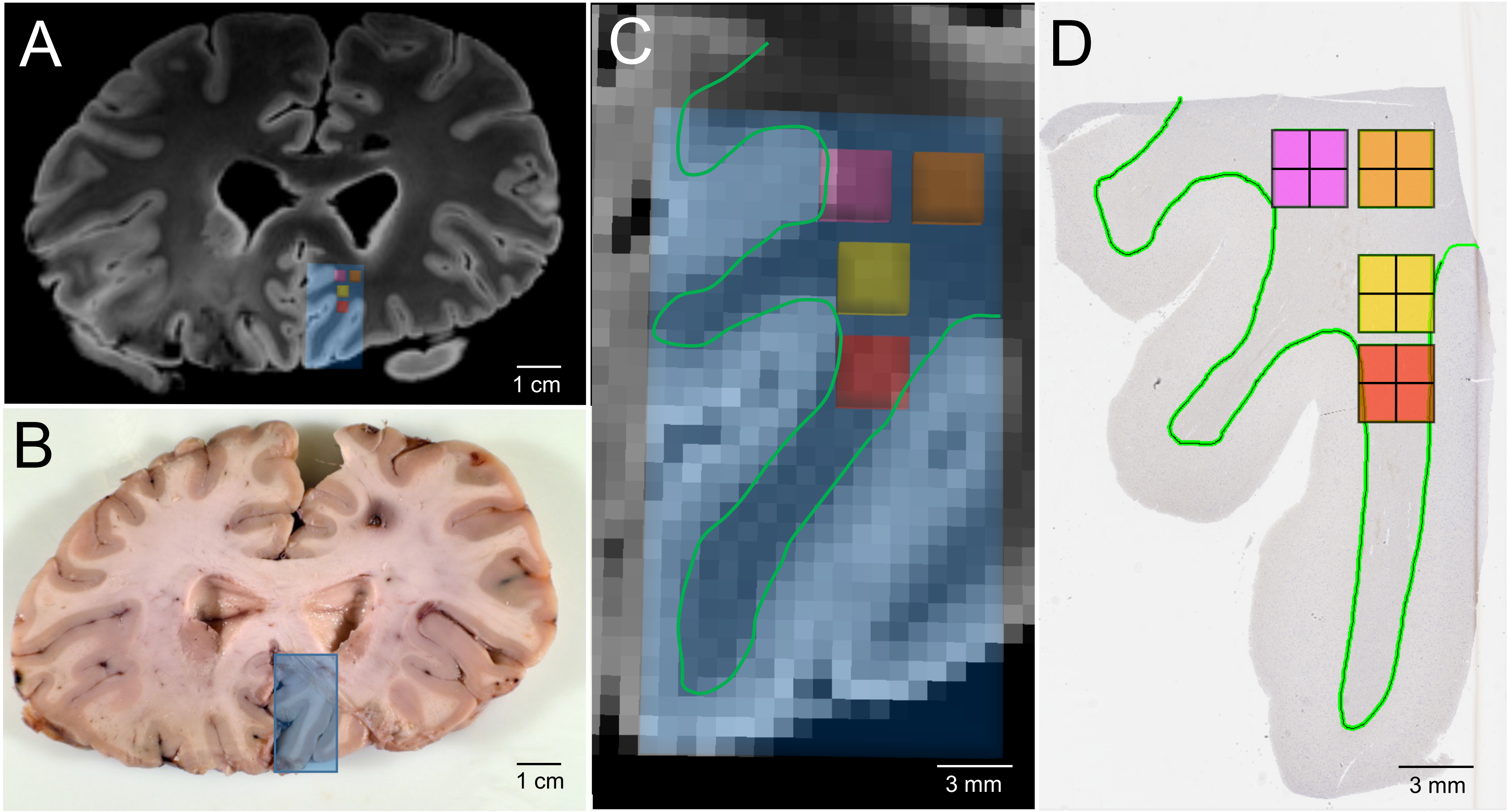


**Supplemental Figure 4. Coregistration of contralateral regions for analysis in *ex vivo* imaging and histologic sections.** A) Coronal section of MRI at the level of the left genu of the corpus callosum. Blue box indicates area sampled for histology. B) Gross pathology of the same region. Blue box indicates the tissue block taken for histology. C) Magnification of area taken for histologic sampling from A. Small pink, orange, yellow and red boxes indicate regions of analysis for tractography, which were co-registered to the histology by mapping their anatomic relationship to the grey-white junction. The green outline in (C) indicates the grey-white junction. D) Image of tissue section with matching color-coded boxes, indicating regions of analysis for histology. A green outline of the grey-white junction is shown in (D) to match the green outline shown in (C).

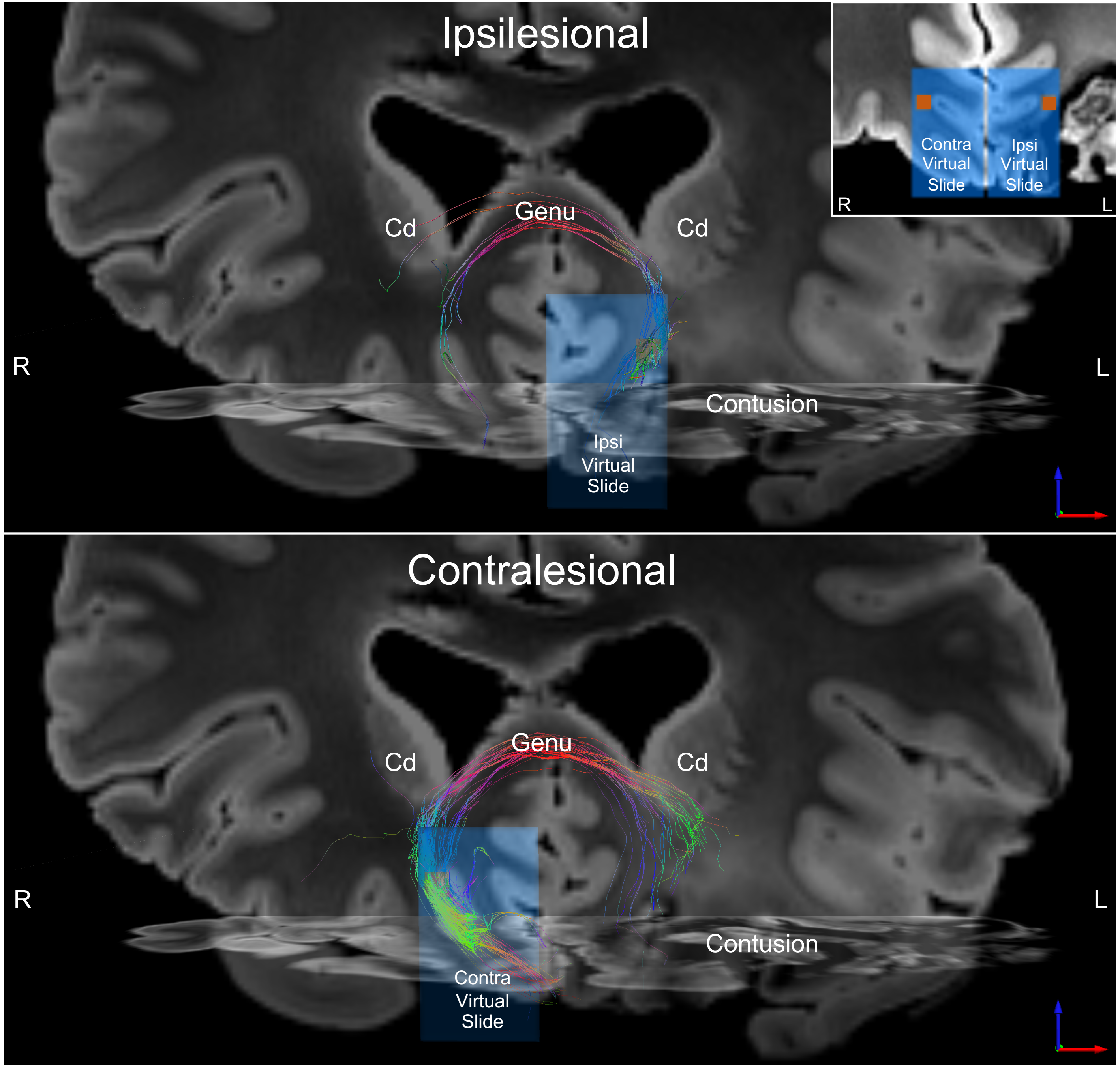


**Supplemental Figure 5. Forceps Minor Tractography Using Ipsilesional and Contralesional Seed Regions.** Anterior views of forceps minor tracts are shown using a region of interest (ROI) ipsilateral to the left frontal contusion and a homologous ROI contralateral to the left frontal contusion. The deterministic tractography data generated from each seed ROI are superimposed on a coronal diffusion-weighted image (DWI) at the level of the genu of the corpus callosum and an axial DWI at the level of the orbitofrontal cortex. Tracts are color-coded by direction (bottom right arrows). In the ipsilesional analysis (top panel), the same ROI shown in Figure 2 was used as a seed to reconstruct the forceps minor tracts. In the contralesional analysis (bottom panel), an ROI with identical size, shape, and anatomic location with respect to the grey-white junction was placed in the right hemisphere. The seed ROIs (each 4x4 voxels) are shown in orange in the top-right inset, superimposed on blue, semitransparent virtual slides. In both tractography analyses, a frontal cortico-cortical bundle was removed to isolate the forceps minor tracts. Visual inspection and quantitative tractography analyses reveal that the contralesional ROI generates more forceps minor tracts (282 tracts) than does the ipsilesional ROI (111 tracts). Furthermore, regardless of where the seed ROI is placed, the forceps minor tracts terminate prematurely in the left hemisphere (i.e. before reaching their left frontal cortical targets) due to the contusion. These results highlight the disruption of forceps minor tracts by the left fontal contusion. Abbreviations: Cd = caudate; L = left; R = right.

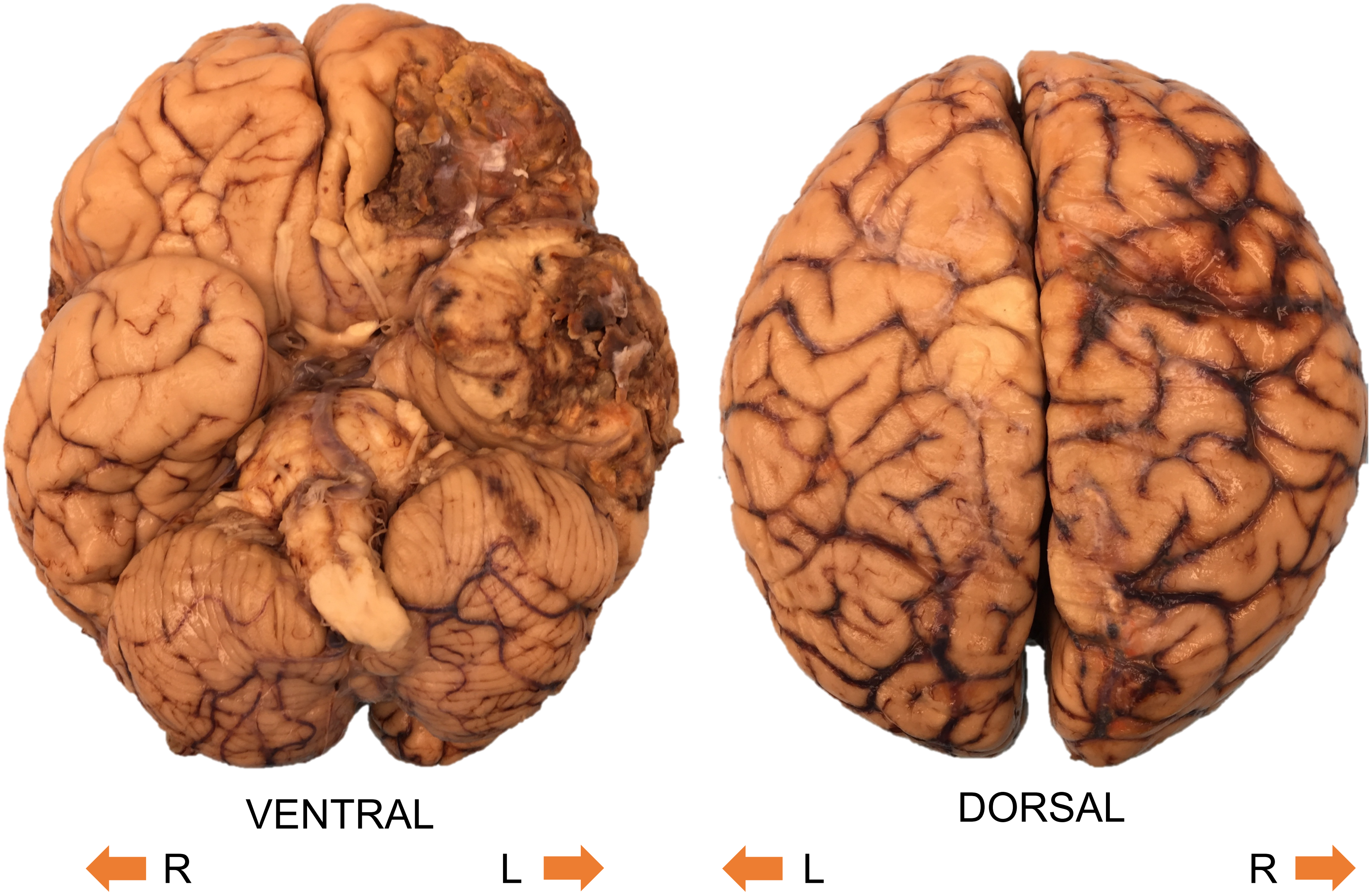


**Supplemental Figure 6**. **Gross Exterior Pathology of the Human Brain Specimen.**
The specimen is shown from a ventral perspective (left) and dorsal perspective (right). Left frontotemporal contusions are seen on the ventral surface of the brain, and sulcal subarachnoid hemorrhage is seen on the right side of the dorsal surface of the brain.

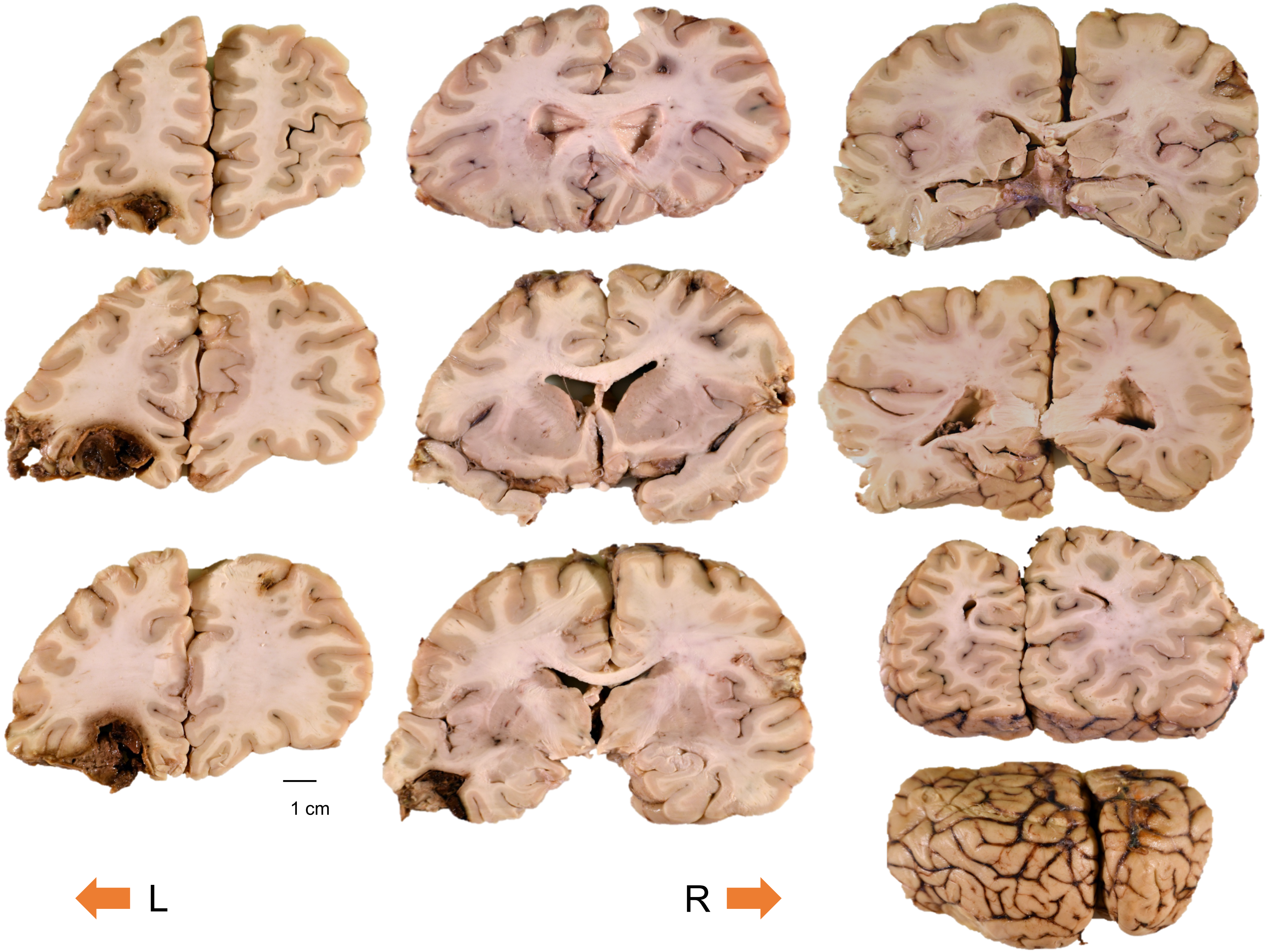


**Supplemental Figure 7. Coronal Slabs of the Cerebral Hemispheres.**
The slabs are displayed from anterior (top left slab) to posterior (bottom right slab).

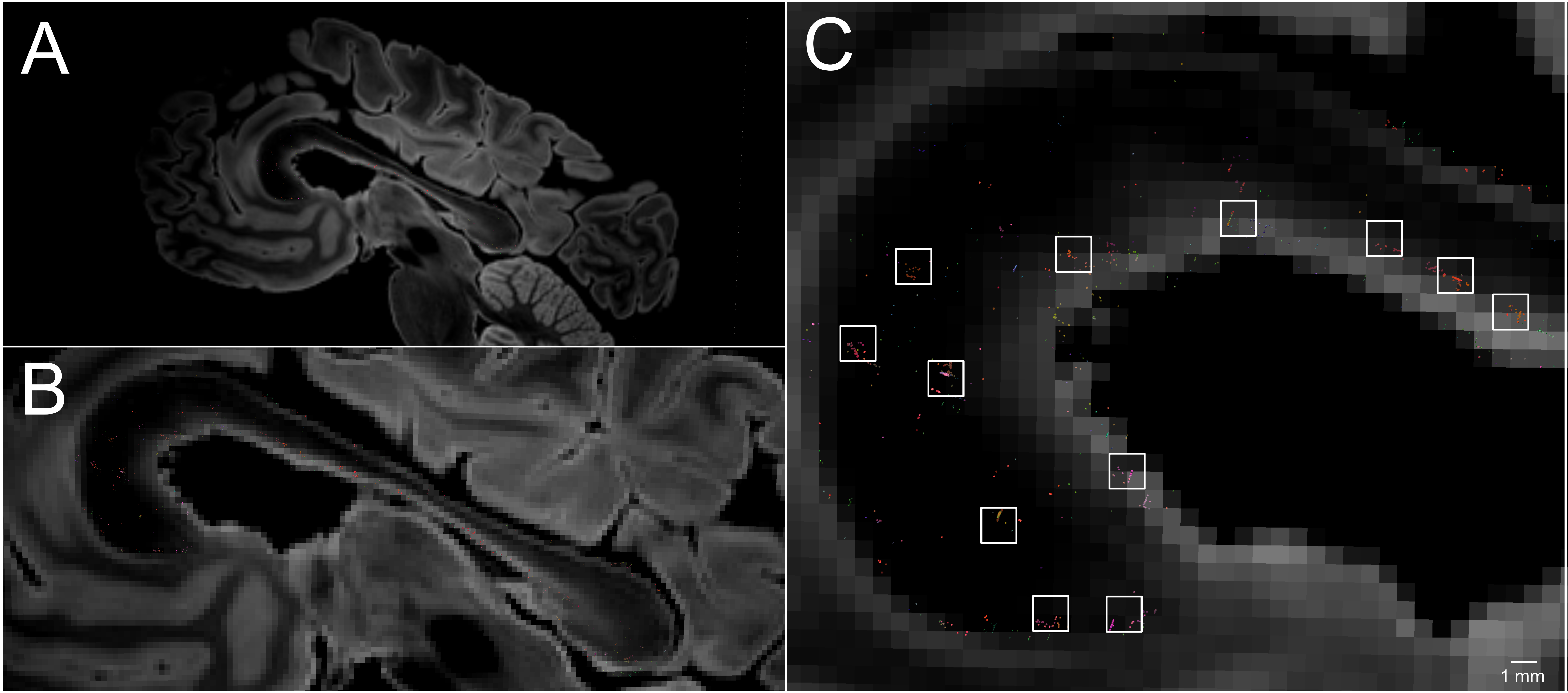


**Supplemental Figure 8. Cluster Analysis of Callosal Tract Disconnections.**
A sagittal diffusion-weighted image (DWI) is shown at the midline from a left lateral perspective in (A). A zoomed view is shown in (B), with endpoints shown for fiber tracts that are disconnected as they pass through a corpus callosum region of interest. Tract endpoints are color-coded by direction (red = medial-lateral; green = antero-posterior; blue = superior-inferior). A high-zoom view is shown in (C). Clusters of disconnected tract end-points are seen in the genu and anterior body of the corpus callosum, located within 2 x 2 voxel (1.5 x 1.5 mm) regions (white squares), indicating that tract endpoints tend to form small clusters rather than being evenly distributed throughout a white matter bundle.

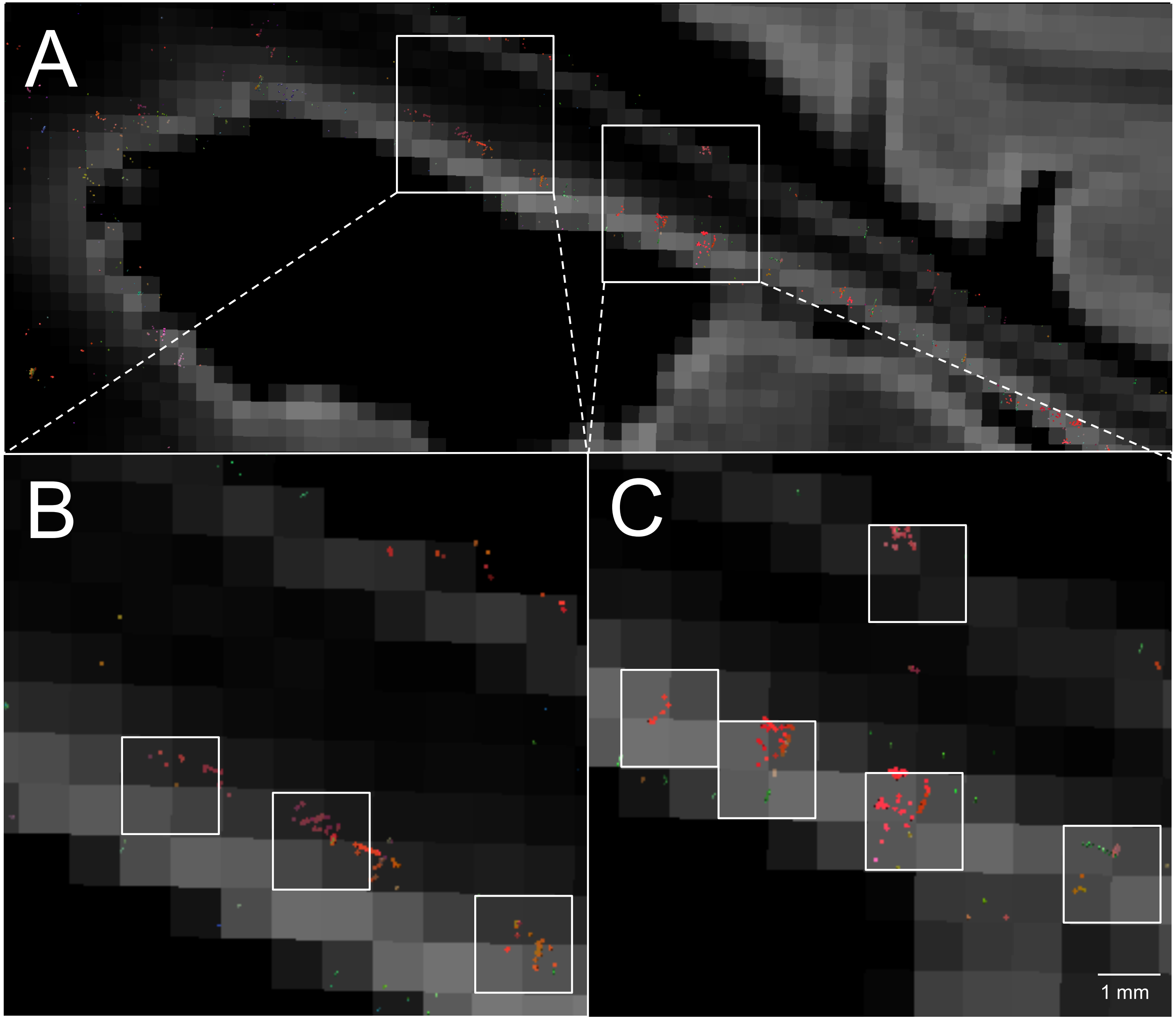


**Supplemental Figure 9. Cluster Analysis of Disconnected Callosal Tracts.**
As in Supplementary Figure 8, a sagittal diffusion-weighted image (DWI) is shown at the midline from a left lateral perspective in (A). Zoomed views of the two regions indicated by white boxes are shown in (B and C), with endpoints shown for fiber tracts that are disconnected as they pass through a corpus callosum region of interest. Tract endpoints are color-coded by direction (red = medial-lateral; green = antero-posterior; blue = superior-inferior). In (B) and (C), clusters of disconnected tract end-points are again seen within 2 x 2 voxel (1.5 x 1.5 mm) regions (white squares) in the mid-body of the corpus callosum, indicating that tract endpoints tend to form small clusters rather than being evenly distributed throughout a white matter bundle.
